# Cyclin D-Cdk4,6 drives cell cycle progression via the retinoblastoma protein’s C-terminal helix

**DOI:** 10.1101/397448

**Authors:** Benjamin R. Topacio, Evgeny Zatulovskiy, Sandra Cristea, Shicong Xie, Carrie S. Tambo, Seth M. Rubin, Julien Sage, Mardo Kõivomägi, Jan M. Skotheim

**Affiliations:** Department of Biology, Stanford University, Stanford, CA 94305, USA; Department of Pediatrics, Stanford University School of Medicine, Stanford, CA 94305, USA; Department of Genetics, Stanford University School of Medicine, Stanford, CA 94305, USA; Department of Chemistry and Biochemistry, University of California, Santa Cruz, CA 95064, USA

**Author notes:** Correspondence (M.K.); (J.M.S.).

**Keywords:** Rb, cyclin, Cdk, E2F, docking, phosphorylation, G1/S, cell cycle regulation

## Abstract

The cyclin-dependent kinases Cdk4 and Cdk6 form complexes with D-type cyclins to drive cell proliferation. A well-known target of cyclin D-Cdk4,6 is the retinoblastoma protein, Rb, which inhibits cell cycle progression until its inactivation by phosphorylation. However, the role of cyclin D-Cdk4,6 phosphorylation of Rb in cell cycle progression is unclear because Rb can be phosphorylated by other cyclin-Cdk complexes and cyclin D-Cdk4,6 complexes have other targets that may drive cell division. Here, we show that cyclin D-Cdk4,6 docks one side of an alpha-helix in the C-terminus of Rb, which is not recognized by cyclins E, A, and B. This helix-based docking mechanism is shared by the p107 and p130 Rb-family members across metazoans. Mutation of the Rb C-terminal helix prevents phosphorylation, promotes G1 arrest, and enhances Rb’s tumor suppressive function. Our work conclusively demonstrates that the cyclin D-Rb interaction drives cell division and defines a new class of cyclin-based docking mechanisms.

## Introduction

The cyclin-dependent kinases Cdk4 and Cdk6 are activated by the D-type cyclins D1, D2, and D3 to drive cell cycle progression from G1 to S phase (Morgan, 1997; Sherr and Roberts, 2004). One important target of Cdk4 and Cdk6 is thought to be the retinoblastoma tumor suppressor protein Rb, which binds and inhibits the activating E2F transcription factors. Rb phosphorylation promotes its dissociation from E2F and thereby drives the expression of E2F-target genes that initiate DNA replication (Bertoli et al., 2013; Dick and Rubin, 2013; Sherr and McCormick, 2002). The importance of Rb phosphorylation and the frequent observation of increased Cdk4 or Cdk6 activity in cancer, has contributed to the consensus model that Cdk4 and Cdk6, activated by their D-type cyclin partners, phosphorylate and inhibit Rb to drive cell cycle progression (Burkhart and Sage, 2008; Lundberg and Weinberg, 1998).

In the current model, cyclin D-Cdk4,6 activity gradually increases until it triggers a positive feedback loop that commits cells to passing the restriction point just prior to the G1/S transition (Merrick et al., 2011; Pardee, 1974; Schwarz et al., 2018). As cells progress through G1, cyclin D-Cdk4,6 would gradually phosphorylate Rb to eventually trigger the onset of E2F-dependent expression of cyclins E and A (Bertoli et al., 2013). Cyclins E and A then bind Cdk1 and Cdk2 to form complexes that continue to phosphorylate Rb (Merrick et al., 2011; Morgan, 2007; Narasimha et al., 2014). In addition, cyclin E and A-dependent Cdk complexes phosphorylate and inhibit the E3 ubiquitin ligase APC/C activating subunit Cdh1, which stabilizes APC/C^Cdh1^ substrates including cyclin A (Di Fiore et al., 2015; Jaspersen et al., 1999; Kramer et al., 2000; Zachariae et al., 1998). The sequential activation of these interconnected positive feedback loops progressively drives commitment to cell division in the face of exposure to anti-proliferative conditions (Cappell et al., 2016; 2018; Pardee, 1974; Schwarz et al., 2018; Yung et al., 2007; Zetterberg and Larsson, 1985).

However, in opposition to the prevailing model of gradually increasing cyclin D-Cdk4,6 activity triggering G1/S, cyclin D levels are nearly constant through G1 (Hitomi and Stacey, 1999). Moreover, Rb is mono-phosphorylated during early-to mid-G1 suggesting that cyclin D-Cdk4,6 activity does not gradually increase through G1 (Narasimha et al., 2014). E2F-dependent transcription increases at the same time that Cdk2 activity increases in late G1. Thus, cyclin D-dependent mono-phosphorylated Rb is still capable of interacting with E2F transcription factors to inhibit transcription. This raises the possibility that Rb inactivation in late G1 is due to hyper-phosphorylation by Cdk2 kinase complexes and that cyclin D-Cdk4,6 promotes the G1/S transition through a different mechanism (Narasimha et al., 2014).

If not Rb, what could be the main target of cyclin D-Cdk4,6 driving cell cycle progression? Possible substrates, whose phosphorylation promotes cell cycle progression, include a mediator of antiproliferative TGF-β signaling Smad3 (Matsuura et al., 2004), an APC/C co-activator Cdh1 (The et al., 2015), and a cell cycle transcription factor FOXM1 (Anders et al., 2011). Furthermore, an additional oncogenic role for Cdk4,6 is likely to be through regulating cellular metabolism, which is consistent with a growing body of literature showing that Cdks directly phosphorylate metabolic enzymes to regulate metabolism in yeast and human cells (Ewald, 2018; Ewald et al., 2016; Salazar-Roa and Malumbres, 2017; Zhao et al., 2016). More specifically, in mammalian cells, cyclin D3-Cdk6 kinase complexes phosphorylate and inactivate the key glycolytic enzymes PFKP and PKM2 to shunt glycolytic intermediates towards NADPH and GSH production, which mitigates ROS accumulation to promote cell survival (Wang et al., 2017). This growing body of literature supports Rb-independent roles for cyclin D-Cdk4,6 in promoting cell proliferation and raises the question as to what is the *in vivo* function of the cyclin D-Rb interaction.

To determine the function of Rb phosphorylation by cyclin D-Cdk4,6, we sought to generate variants of Rb that could no longer interact with cyclin D-Cdk4,6 but maintained all the other interactions of cyclin D-Cdk4,6. The specificity of substrate binding and phosphorylation by cyclin-Cdk complexes is generally determined by the ability of the cyclin to recognize docking sites on substrates (Morgan, 2007). Previously identified docking sites on substrates are short linear amino acid motifs. In budding yeast, the G1 cyclin Cln2 recognizes an LP docking motif and the S Phase cyclins Clb5 and Clb3 recognize substrates with RxL docking motifs (Bhaduri and Pryciak, 2011; Cross and Jacobson, 2000; Kõivomägi et al., 2011; Loog and Morgan, 2005). In animal cells, cyclin A-Cdk2 and cyclin E-Cdk2 complexes also utilize RxL-based docking through their *α*1 helix hydrophobic patches to bind and phosphorylate substrate proteins including Rb, p107, p27, and Cdc6 (Adams et al., 1999; Hirschi et al., 2010; Russo et al., 1996; Schulman et al., 1998; Takeda et al., 2001; Wohlschlegel et al., 2001). While cyclin D likely has a hydrophobic patch on its *α*1 helix similar to cyclins E and A, it has not been shown if this patch also recognizes RxL motifs. Cyclin D has an N-terminal LxCxE motif, which binds Rb’s LxCxE cleft (Dick and Rubin, 2013; Dowdy et al., 1993). However, mutation of the LxCxE cleft had only a modest effect *in vivo* suggesting that either the cyclin D-Rb interaction is of limited importance or that there exists additional important docking interactions. Supporting the existence of an additional cyclin D-Rb docking mechanism, truncation of the Rb C-terminus disrupted phosphorylation by cyclin D1-Cdk4 *in vitro,* and this truncated Rb variant slowed division and inhibited E2F-dependent gene expression in cultured cells (Gorges et al., 2008; Pan et al., 2001; Wallace and Ball, 2004).

In this study, we analyzed the docking interactions between Rb and cyclin D-Cdk4,6 complexes. We found that cyclin D-Cdk4,6 targets the Rb family of proteins (Rb, p107, and p130) for phosphorylation primarily by docking a C-terminal alpha-helix. Importantly, this Rb C-terminal helix is not recognized by the other major cell cycle cyclin-Cdk complexes cyclin E-Cdk2, cyclin A-Cdk2, and cyclin B-Cdk1. Thus, mutation of this helix disrupts cyclin D’s ability to phosphorylate Rb, but maintains Rb regulation by other cyclin-Cdk complexes. Disruption of helix-based docking reduced Rb phosphorylation, induced G1 cell cycle arrest in cell lines, and slowed tumor growth *in vivo*. Taken together, our results show that cyclin D-Cdk4,6 phosphorylates and inhibits Rb and that this interaction is a major driver of cell proliferation.

## Results

### RxL and LxCxE based docking mutations broadly affect cyclin-Cdk complexes

To determine the function of Rb phosphorylation by cyclin D-Cdk4,6, we sought to generate an Rb protein variant that does not interact with cyclin D-Cdk4,6, but does interact with other cyclin-Cdk complexes (Figure 1A-B). To identify specific cyclin D-Rb interactions, we performed *in vitro* kinase assays on Rb protein variants with a panel of purified cyclin-Cdk complexes (Schwarz et al., 2018) (Figure S1). In contrast to other work on Rb phosphorylation working with purified Rb fragments, we were able to purify full-length Rb protein from bacteria. In these kinase assay experiments, mutation of Rb residues required for docking interactions would then manifest as reduced kinase activity towards Rb. Compared with other cyclin-Cdk complexes, cyclin D-Cdk4,6 exhibited no detectable kinase activity towards the model Cdk substrate histone H1 (Kato et al., 1993; Matsushime et al., 1994) but was capable of phosphorylating Rb (Figure 1C-E).

**Figure 1.**
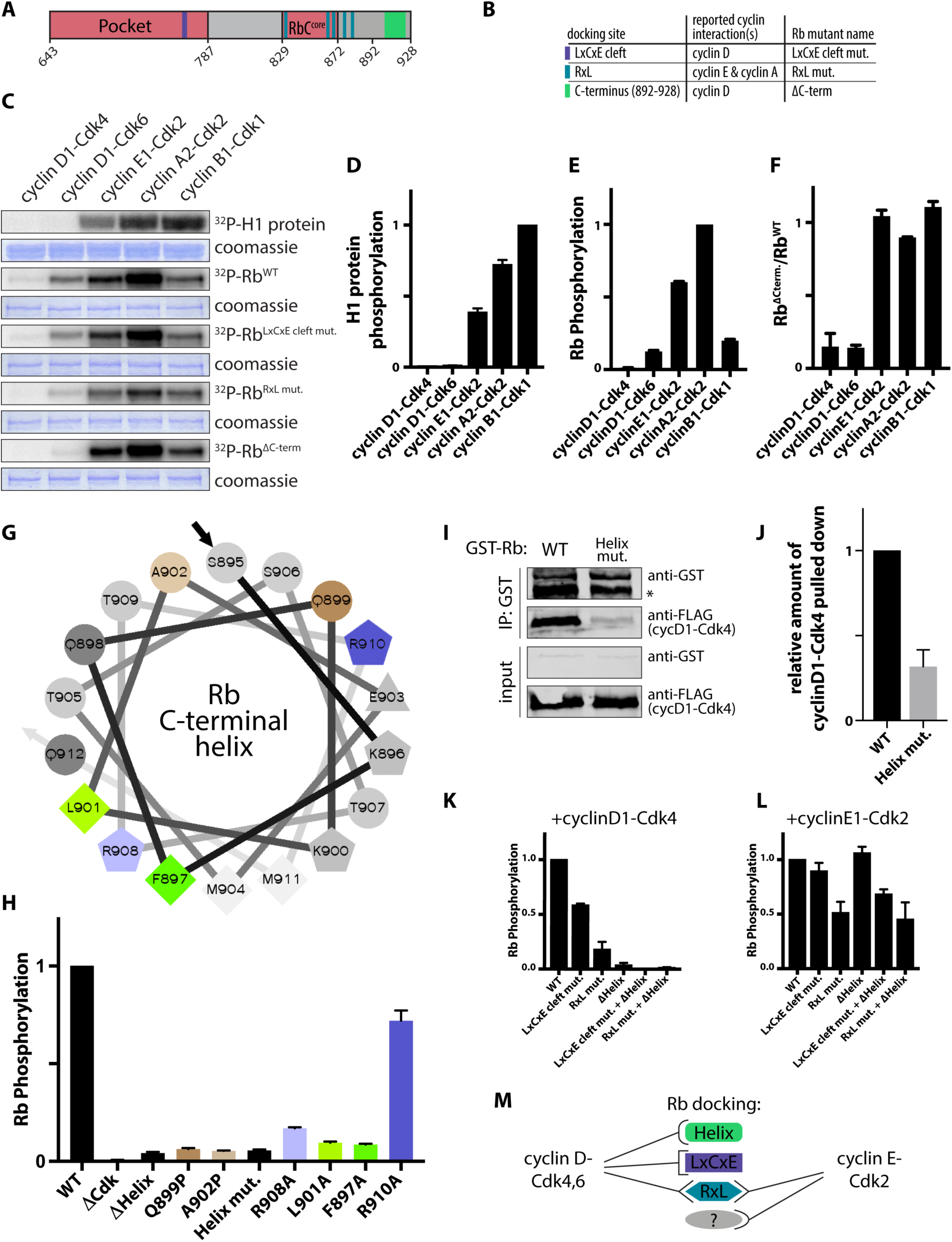
Cyclin D-Cdk4,6 complexes target Rb for phosphorylation by docking a C-terminal helix. (A and B) Summary of cyclin-Cdk docking sites on Rb (amino acids 643 to 928) (A) and names of the docking site mutants (B). (C) *In vitro* kinase assays using the denoted cyclin-Cdk complexes and histone H1 or variants of Rb. Rb^RxL mut.^ lacks all C-terminal RxL sequences that dock cyclin hydrophobic patches. Rb^LxCxE cleft mut.^ lacks the LxCxE docking cleft. Rb^ΔC-term^ lacks the C-terminal amino acids 892-928. The coomassie stained gel showing equal amounts of substrate used in each reaction is placed below each autoradiograph. (D and E) Quantification of H1 kinase assays (D) and Rb kinase assays (E) using the denoted cyclin-Cdk complexes. (F) The ratio of kinase activity of the denoted cyclin-Cdk complexes towards Rb and Rb^ΔC-term^. (G) Helical wheel projection of the predicted Rb C-terminal helix. Shapes denote residues with similar characteristics. Colored amino acids correspond to alanine or proline substitutions in (H). (H) *In vitro* kinase assays of Rb protein variants by cyclin D1-Cdk4. ΔCdk denotes Rb lacking the 15 Cdk phosphorylation sites. ΔHelix denotes Rb lacking the C-terminal helix amino acids 895-915. Helix mut. denotes an Rb variant where the predicted docking interface residues F897, L901, and R908 are substituted with alanines. (I) Western blot of a GST-pulldown binding assay. The GST-tagged Rb bait proteins were incubated with 3X FLAG-tagged cyclin D1 prey proteins. *denotes a degradation product below full-length GST-Rb. (J) Quantification cyclin D1-Cdk4 pulled down in (F). (K and L) Quantification of *in vitro* kinase assays of the denoted Rb variant with (K) cyclin D1-Cdk4 or (L) cyclin E1-Cdk2. Data shown are mean ± SEM. (D-E) and (H), *n* = 2; (J), *n* = 3; (K and L), *n* = 3.

To test the effect of mutating known cyclin D-Rb docking interactions, we first removed the LxCxE binding cleft in the Rb pocket domain that interacts with proteins containing the LxCxE motif, such as viral oncoproteins and cyclin D (Dick et al., 2000; Dowdy et al., 1993; Markey et al., 2007). This resulted in only a 1.7±0.3-fold reduction in phosphorylation by cyclin D kinase complexes (Figure 1C and S1J), consistent with reports that the LxCxE-docking interaction alone is weak and its removal had only a modest effect in cells (Dick et al., 2000; Guiley et al., 2015). Moreover, we observed that previously reported LxCxE cleft mutations similarly affected cyclin E-, A-, and B-dependent phosphorylation of Rb *in vitro* and therefore may not be specific for cyclin D (Figure 1C and S1J).

Next, we tested the effect of mutating the RxL motifs on Rb that were shown to interact with the S phase cyclins E and A (Adams et al., 1999; Hirschi et al., 2010; Schulman et al., 1998). We generated an Rb variant protein lacking all 5 RxL sequences in its unstructured regions. Compared to wild type Rb, this RxL variant Rb was phosphorylated 2.0±0.1-fold less by cyclin E-Cdk2 and 2.9±0.2-fold less by cyclin A-Cdk2 (Figure 1C and S1J). This implies that while cyclins E and A do use RxL docking as previously suggested, this is not the only mechanism they use to identify their substrates. Cyclin B-Cdk1 phosphorylation of Rb was unchanged by mutating the RxL motifs, suggesting that cyclin B does not use its hydrophobic patch to phosphorylate Rb. While it has not been studied extensively, cyclin D has an *α*1 helix hydrophobic patch as cyclins E, A and B, but comprised of different residues (Morgan, 2007). We observed that the Rb variant lacking RxL motifs was phosphorylated 4.1±0.4-fold less by cyclin D-Cdk4,6, suggesting that cyclin D recognizes RxL motifs as other cyclins (Figure 1C and S1J). While the RxL docking is clearly important for cyclin D, it is shared with other cyclins so that its mutation would not specifically disrupt the cyclin D-Rb interaction.

### Cyclin D-Cdk4,6 complexes target Rb for phosphorylation by docking a C-terminal helix

To generate an Rb protein variant that does not interact with cyclin D-Cdk4,6, but does interact with other cyclin-Cdk complexes we next examined the effect of truncating the final 37 amino acids of Rb. Truncation of the Rb C-terminus, which included these 37 amino acids, was previously shown to reduce Rb phosphorylation *in vitro* and promote Rb’s ability to arrest cells in G1 (Gorges et al., 2008; Pan et al., 2001; Wallace and Ball, 2004). C-terminal truncation of Rb reduced phosphorylation by cyclin D1-Cdk4,6 20±10-fold (Figure 1C and S1J) and increased the Michaelis-Menten constant K_M_ beyond our measurement limit of ∼5µM (Figure S2G-H), indicating the presence of a docking interaction in the final 37 amino acids of the Rb C-terminus. Importantly, this C-terminal truncation is specific for cyclin D-Cdk4,6 complexes and does not affect the phosphorylation of Rb by cyclin E-Cdk2, cyclin A-Cdk2, and cyclin B-Cdk1 complexes (Figure 1C and 1F).

We next sought to determine how the Rb C-terminus promotes phosphorylation by cyclin D-Cdk4,6. Cyclin substrate docking has previously been shown to arise from short linear motifs of a few amino acids in intrinsically disordered regions on the target proteins, a common mechanism for evolution of protein-protein interactions (Bloom and Cross, 2007; Davey et al., 2015). However, such a short linear motif model is unlikely to explain the interaction of cyclin D with the Rb C-terminus because *in vitro* Rb phosphorylation by cyclin D-Cdk4 is affected by mutations over a large range of amino acids (Wallace and Ball, 2004). While the Rb C-terminus is intrinsically disordered, it is known to adopt structure when bound to other proteins (Rubin et al., 2005). We therefore examined the far C-terminal Rb amino acids for potential secondary structure and found a region with alpha-helix propensity spanning 21 amino acids (Rb 895-915; Figure 1G and S3). Deletion of this potential helix (ΔHelix) or disruption of this helix by proline substitution (Q899P or A902P) drastically reduced Rb phosphorylation by cyclin D-Cdk4,6, which was comparable to the phosphorylation of an Rb variant (ΔCdk) lacking all 15 Cdk phosphorylation sites (Figure 1G-H).

The Rb C-terminal helix is predicted to have one face composed primarily of hydrophobic residues that could interact with cyclin D-Cdk4,6 (Figure 1G). This potential interface contains two hydrophobic residues, F897 and L901, and a positively charged residue, R908, that line up on one side of the helix. Alanine substitution of these three amino acids, denoted as Rb^Helix mut.^, disrupts phosphorylation of Rb by cyclin D-Cdk4,6 (Figure 1G-H and Figure S2A-H). Since alanines have high helical propensity (Nick Pace and Martin Scholtz, 1998) these mutations likely disrupt the recognition of Rb by cyclin D-Cdk4,6 without disrupting the C-terminal helix (Figure S3B). In contrast, substitution of an arginine amino acid on the opposite face of the helix (R901A) has only a modest effect on phosphorylation (Figure 1G-H). To test if the predicted interface mutations affect cyclin D-Cdk4,6 binding to Rb, we performed GST-pulldown assays. Further supporting the helix-docking model, mutation of the helix interface disrupted the binding of Rb and cyclin D-Cdk4,6 (Figure 1I-J).

Finally, we examined the effect of combining mutations on Rb (Figure 1K-L and S2I). LxCxE, RxL, and helix mutations were additive, suggesting that these docking sites interact with different parts of cyclin D. Taken together, our analysis shows that there is a diversity of poorly understood docking mechanisms that are distinct for each cyclin (Figure 1M).

### Cyclin D1-Cdk4,6 phosphorylates synthetic substrates fused to the Rb C-terminal helix

To further test the helix-based docking model, we sought to confer increased cyclin D activity to poor substrates such as an Rb peptide containing a single Cdk consensus phosphorylation site (Grafström et al., 1999; Kitagawa et al., 1996). To do this, we generated a fusion protein containing a GST-tag, Rb amino acids 775-790 that contain a single Cdk site, a Gly-Ser linker, and the Rb C-terminal helix (Figure 2A, S4, and Table S1). Fusing the helix to this previously poor substrate led to a 13.9±0.1-fold increase in phosphorylation by cyclin D1-Cdk6, but did not affect phosphorylation by cyclin E1-Cdk2 (Figure 2B-C). We also observed a similar increase in activity towards fusion proteins containing an Cdk site on a peptide derived from Histone H1 (Matsushime et al., 1994) or containing Rb amino acids 790-805, which has a single Cdk site (Figure S4E-F). Cyclin D1-Cdk6 did not phosphorylate these synthetic substrates if they were fused to the Rb helix with the predicted interface residues mutated to alanine (Figure 2B, S4A, and S4D-E). To test if orientation of these interface residues was important, we generated a synthetic substrate with a single Cdk site that was fused to a helix composed of the reversed amino acid sequence of the Rb C-terminal helix. This reversed helix substrate was poorly phosphorylated by cyclin D1-Cdk6 implying that the orientation of the interface is important, in addition to the polarity and acid-base properties of the interface residues (Figure 2A-B, S4A, and S4D). As predicted from our analysis of the full-length Rb protein, fusion of the Rb C-terminal helix did not affect the phosphorylation of these synthetic substrates by cyclin E1-Cdk2 (Figure 2C, S4B, S4D, and S4F). Taken together, these data show that the Rb C-terminal helix is sufficient to direct cyclin D-Cdk4,6 complexes to phosphorylate substrates, regardless of any intrinsic preference for a particular Cdk site. Moreover, the Rb C-terminal helix is specific for cyclin D-Cdk4,6 complexes (Figure 2D-E).

**Figure 2.**
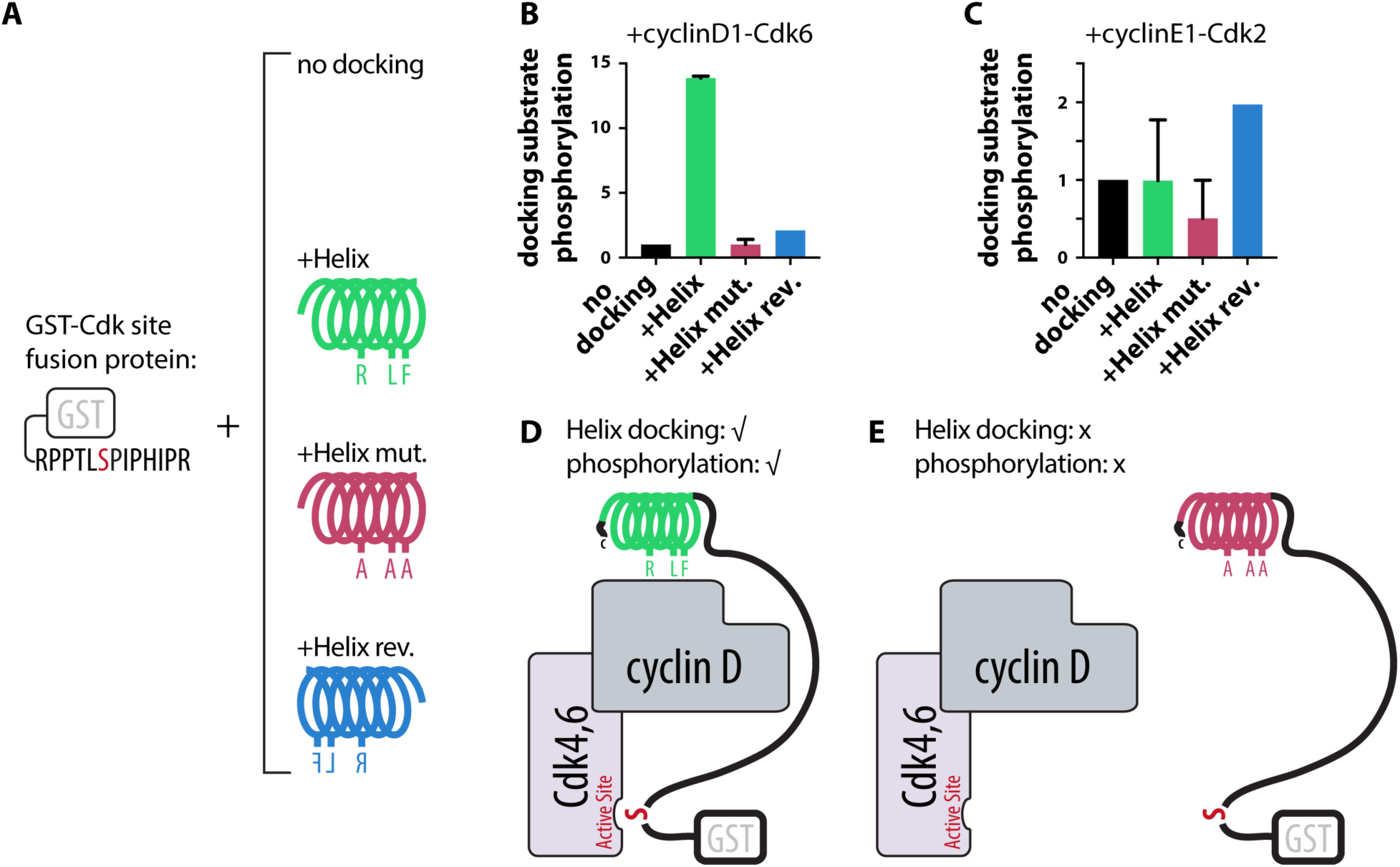
Cyclin D1-Cdk4,6 phosphorylates synthetic substrates fused to the Rb C-terminal helix. (A) Schematic of the engineered GST-Cdk phosphorylation site fusion protein containing a GST tag and the Rb amino acids 775-787 containing a single Cdk site fused to either no docking site, the Rb C-terminal helix docking site (+Helix), the Rb C-terminal helix docking site with the three interface residues substituted with alanines (+Helix mut.), or the Rb C-terminal helix docking site in reverse (+Helix rev.). (B and C) *In vitro* kinase assays of the indicated engineered GST-Cdk phosphorylation site fusion protein by the indicated kinase. (D and E) Schematic of docking interaction between cyclin D-Cdk4,6 complexes and the engineered GST-Cdk phosphorylation site fused to (D) the Rb C-terminal helix docking site, or (E) its mutated version. Data shown are mean ± SEM. (B and C), *n* = 3.

### D-type cyclins recognize the Rb C-terminal helix

To determine the subunit of the cyclin D-Cdk4,6 complex that interacts with the Rb C-terminal helix, we first generated all six possible complexes formed by cyclins D1, D2 and D3 with either Cdk4 or Cdk6. All six possible cyclin D-Cdk4,6 complexes target Rb for phosphorylation through helix-based docking (Figure 3A). Since all three D-type cyclins have a hydrophobic patch, we sought to determine what this patch recognized. To do this, we purified Cdk6 in complex with a cyclin D1 variant in which the residues that form the hydrophobic patch were substituted with alanines (cyclin D1^HPmut.^). Compared to wild type cyclin D1-Cdk6, cyclin D1^HPmut.^-Cdk6 exhibited weaker kinase activity towards Rb consistent with a function for the hydrophobic patch (Figure 3B). However, cyclin D1^HPmut.^-Cdk6 did not phosphorylate an Rb variant where the C-terminal helix interface residues were substituted with alanines. This implies that the hydrophobic patch is not responsible for recognizing the Rb C-terminal helix (Figure 3B).

**Figure 3.**
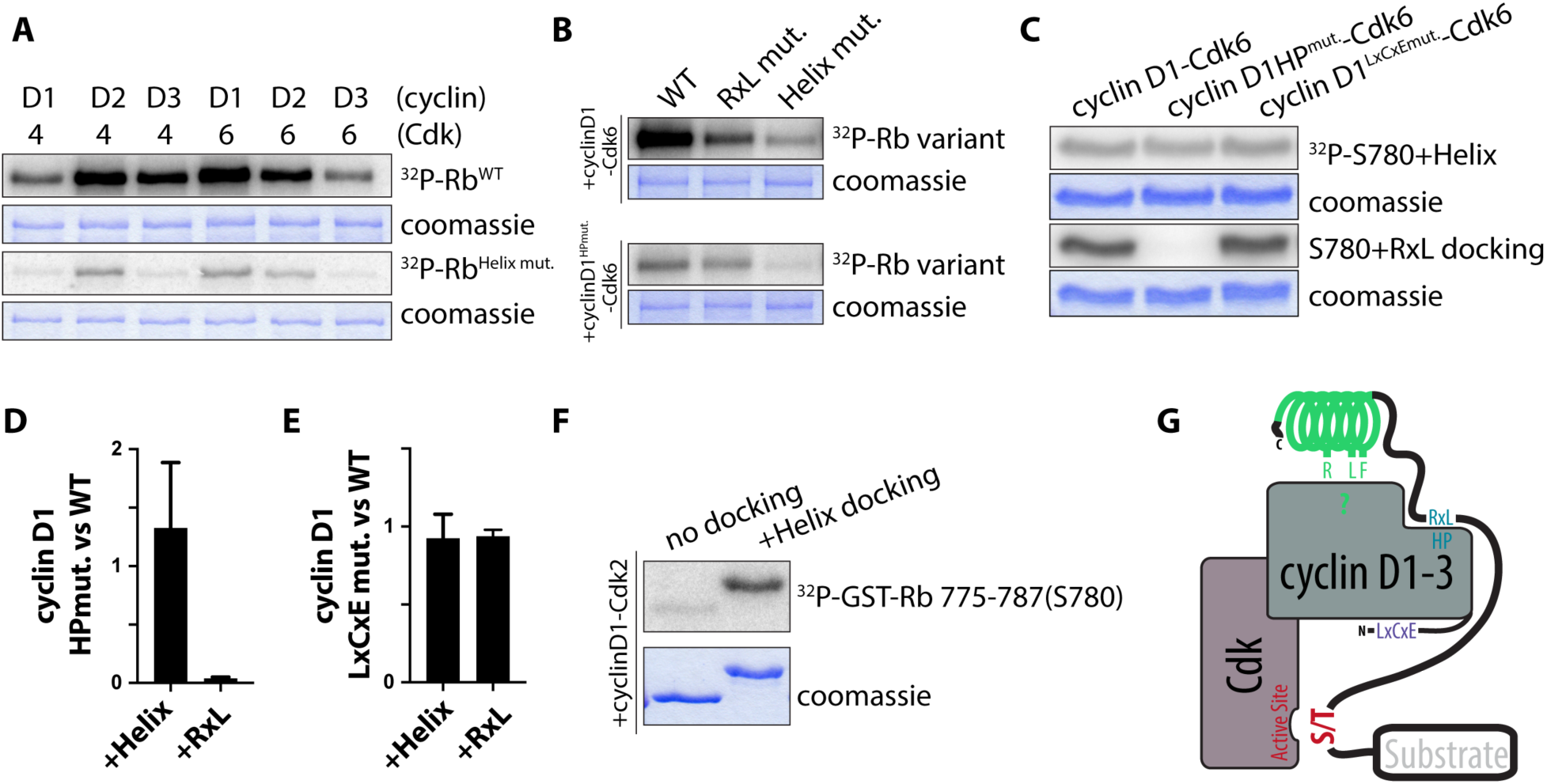
D-type cyclins recognize the Rb C-terminal helix. (A) *In vitro* kinase assays with all six possible cyclin D-Cdk4,6 complexes and the denoted Rb variants. Fold changes ± SEM of wild-type Rb to Helix mutant Rb phosphorylation are: cyclin D1-Cdk4 = 30±10; cyclin D2-Cdk4 = 19±3; cyclin D3-Cdk4 = 90±20; cyclin D1-Cdk6 = 17±6; cyclin D2-Cdk6 = 19±8; cyclin D3-Cdk6 = 30±20. (B) *In vitro* kinase assays of the denoted Rb variant with cyclin D1-Cdk6 or cyclin D1^HPmut.^-Cdk6. (C) *In vitro* kinase assays using the denoted cyclin D1-Cdk6 complexes with GST-Rb775-787(S780)+Helix docking or GST-Rb775-787(S780)+RxL docking. Cyclin D1^HPmut.^ denotes mutation of the hydrophobic patch on cyclin D1. Cyclin D1^LxCxE mut.^ denotes mutation of the LxCxE motif on cyclin D1. (D and E) Quantification of the phosphorylation of the engineered GST-Cdk phosphorylation site fusion proteins containing helix docking from Rb (+Helix) or RxL docking from Cdc6 (+RxL) by wild-type cyclin D1-Cdk6 relative to (D) cyclin D1^HPmut.^-Cdk6 or (E) relative to cyclin D1^LxCxEmut.^-Cdk6. (F) *In vitro* kinase assays using cyclin D1 fused to Cdk2. GST-Rb775-787(S780) without docking or GST-Rb775-787(S780)+Helix docking were used as substrates. (G) Schematic of docking interaction between cyclin D1, D2, and D3 complexes and their substrates. The coomassie stained gels showing equal amounts of substrate used in each reaction are placed below each autoradiograph. Data shown are mean ± SEM. (A and B) representative experiments shown (out of two independent experiments); (D and E) *n* = 3; (F) representative experiments shown (out of two independent experiments).

To test if cyclin D’s hydrophobic patch docks RxL sequences like cyclin A and E (Adams et al., 1999; Hirschi et al., 2010; Schulman et al., 1998; Takeda et al., 2001), we compared the ability of cyclin D1-Cdk6 and cyclin D1^HPmut.^-Cdk6 to phosphorylate synthetic substrates containing an RxL motif. Mutation of the cyclin D hydrophobic patch removed the ability of cyclin D1-Cdk6 to phosphorylate a fusion protein containing a GST tag, the Rb amino acids 775-790 containing a single Cdk site, and a peptide containing an RxL sequence derived from the known Cdk substrate Cdc6 (Takeda et al., 2001) (Figure 3C-D and S4C). However, the cyclin D hydrophobic patch mutation did not affect the ability of cyclin D1-Cdk6 to phosphorylate a fusion protein where the RxL docking sequence was replaced by the Rb C-terminal helix (Figure 3C-D). As expected, mutation of the LxCxE sequence in cyclin D did not affect phosphorylation of any of these substrates (Figure 3C and 3E). Taken together, these data show that the hydrophobic patch on cyclin D recognizes linear RxL sequences including those on Rb.

That cyclin D’s hydrophobic recognized RxL sequences but not the Rb C-terminal helix raised the question if cyclin D or Cdk4,6 was responsible for helix docking. To test if cyclin D was responsible for helix-based docking, we purified cyclin D1-Cdk2 complexes and observed that cyclin D1-Cdk2 complexes phosphorylate synthetic substrates containing the C-terminal helix (Figure 3F). Since cyclin E1-Cdk2 complexes does not use helix docking, our experiments imply that the helix-docking site likely lies on cyclin D rather than Cdk4,6 since the cyclin D1-Cdk2 fusion can use helix-based docking. Thus, the hydrophobic patch of cyclin D likely recognizes RxL sequences, while a different region on cyclin D is likely responsible for recognizing the Rb C-terminal helix (Figure 3G).

### A C-terminal docking helix is present across the metazoan Rb protein family

We next examined if other cyclin D-Cdk4,6 substrates used helix-based docking. As few Cdk4,6 substrates are known (Anders et al., 2011; Malumbres and Barbacid, 2005), we initially examined the Rb family members, p107 and p130, which are similar in sequence and function (Dick and Rubin, 2013). Based on their sequence, we predict that all human and mouse Rb family proteins have C-terminal helices, which we found targeted them for phosphorylation by cyclin D-Cdk4,6 *in vitro* (Figure 4A-D and Figure S5). Alanine substitutions at what we predict to be the docking interface for p107 and p130 also disrupted phosphorylation by cyclin D1-Cdk4 (Figure 4A-D). We observed a similar requirement for helix-based docking with mouse Rb phosphorylation (Figure S5K-L). The presence of helical docking across the Rb family led us to search for the conserved sequence motif, which we identified by aligning 682 C-terminal sequences of metazoan Rb family members (Medina et al., 2016) and generating a consensus sequence motif (Figure 4E). We did not find such C-terminal helices in Rb sequences outside metazoans, suggesting that C-terminal helical docking is a metazoan innovation (Figure 4F). Taken together, our results suggest that cyclin D-Cdk4,6 targets the Rb family through a similar mechanism across metazoans. A motif search through the human proteome looking for such docking helices on proteins containing Cdk consensus phosphorylation sites suggests as many as 70 proteins may use this mechanism (Table S2).

**Figure 4.**
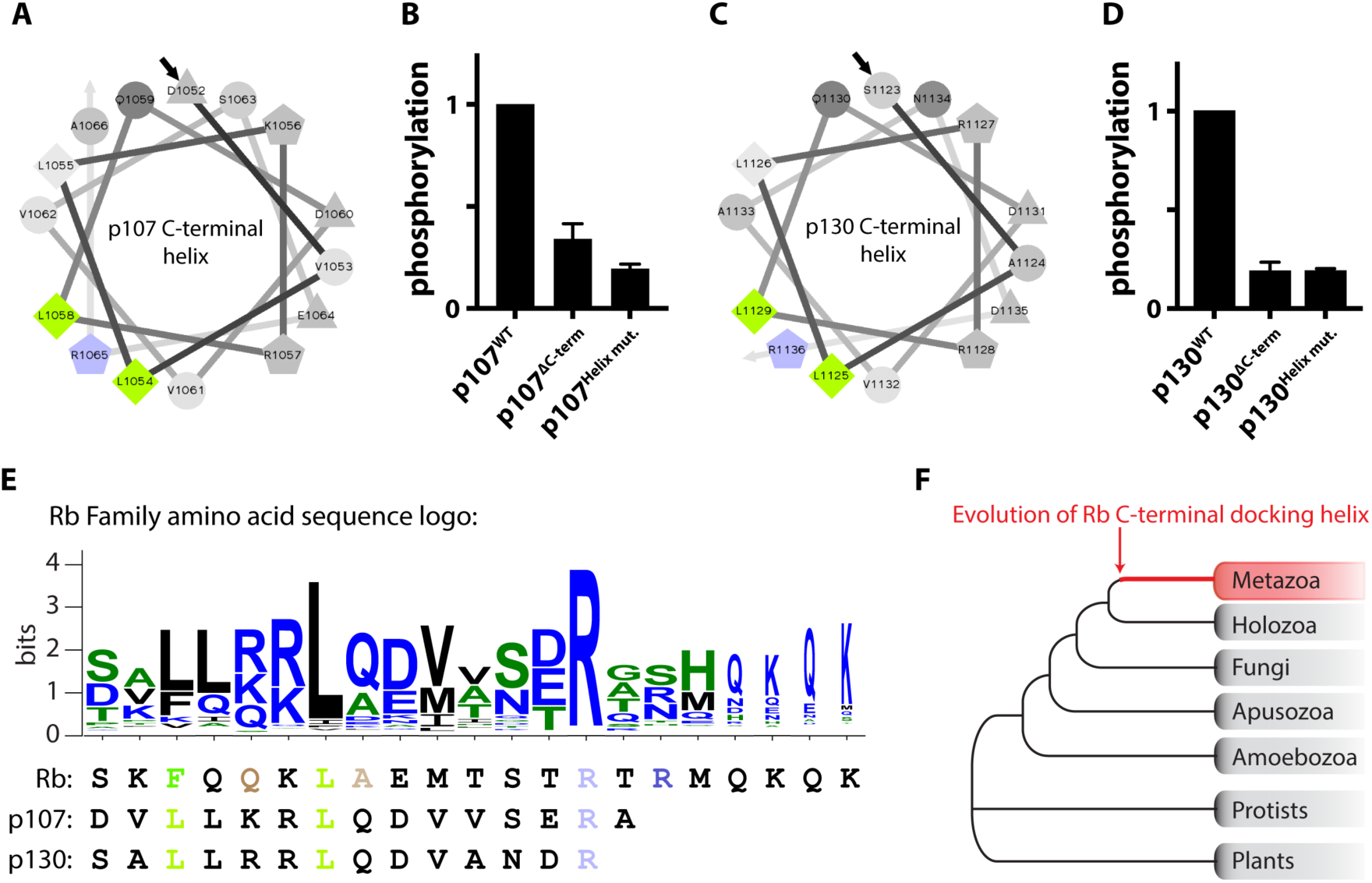
A C-terminal docking helix is present across the metazoan Rb protein family. (A-D) Helical wheel projection of the predicted C-terminal helix for (A) human p107 and (C) p130. *In vitro* kinase assays of human (B) p107 and (D) p130 variants by cyclin D1-Cdk4. ΔC-term denotes truncating p107 at amino acid position 1014 and p130 at amino acid position 1122. Helix mut. denotes alanine substitution of the predicted docking interface residues indicated in blue and green in (A) and (C). (E) C-terminal helix sequence motif generated using 682 sequences of metazoan Rb family members. In the logo, blue denotes hydrophilic residues, green denotes neutral amino acids, and black denotes hydrophobic amino acids. Below the logo, C-terminal helix residues for human Rb, p107, and p130 are aligned. The C-terminal helix residues that were tested for importance to cyclin D-docking are colored according to Figure 1G, (A), and (C). (F) A C-terminal Rb helix was not found outside metazoan genome sequences. Data shown are mean ± SEM. (B and D) *n* = 2.

### The cyclin D-Rb interaction promotes the G1/S transition, Rb dissociation from chromatin, and E2F1 activation

The identification of the C-terminal docking helix on Rb allowed us to test the function of Rb phosphorylation by cyclin D-Cdk4,6 in cell cycle control. This is because mutation of Rb’s C-terminal helix disrupts phosphorylation by cyclin D-Cdk4,6, but not other cell cycle-dependent cyclin-Cdk complexes so that the introduction of Rb helix mutations specifically test the function of the cyclin D-Rb interaction.

To test the function of the cyclin D-Rb interaction, we examined immortalized human mammary epithelial cells (HMECs) expressing doxycycline-inducible Rb variant proteins fused to Clover fluorescent protein and 3FLAG affinity tags (Figure 5A-B and S6A-B). We chose to examine HMECs because they are a non-transformed cell line previously used to study cell growth and proliferation (Sack et al., 2018). Expression of exogenous wild-type Rb in HMECs had a minor effect on cell cycle progression and cell size, whereas expression of Rb^Helix mut.^, which lacks helix-based docking, increased the fraction of cells in G1 from 50% to 75%, and increased cell size by 55% (Figure 5C-D). Expression of Rb^Helix mut^ had a similar effect to exposing cells to 500nM palbociclib, a Cdk4,6 inhibitor (Figure 5C). Expression of the double mutant Rb^LxCxE cleft+Helix mut.^ protein, which lacks LxCxE- and helix-based docking mechanisms, resulted in a dramatic G1 cell cycle arrest and a cell size increase similar to the effect of expressing an Rb^ΔCdk^ protein lacking all 15 Cdk phosphorylation sites (Figure 5C-D). The cell size and cell cycle phase phenotypes correlated with the amount of chromatin-bound Clover-3FLAG-Rb in G1 phase cells for each Rb variant (Figure 5E). More specifically, following a low salt wash (Håland et al., 2015; Lundberg and Weinberg, 1998), there was 11.5-fold more Rb^ΔCdk^ than wild-type Rb that remained bound to chromatin in G1 in a representative experiment. These effects were not due to differences in Rb protein expression as measured using a quantitative immunoblot (Figure S6A-B).

**Figure 5.**
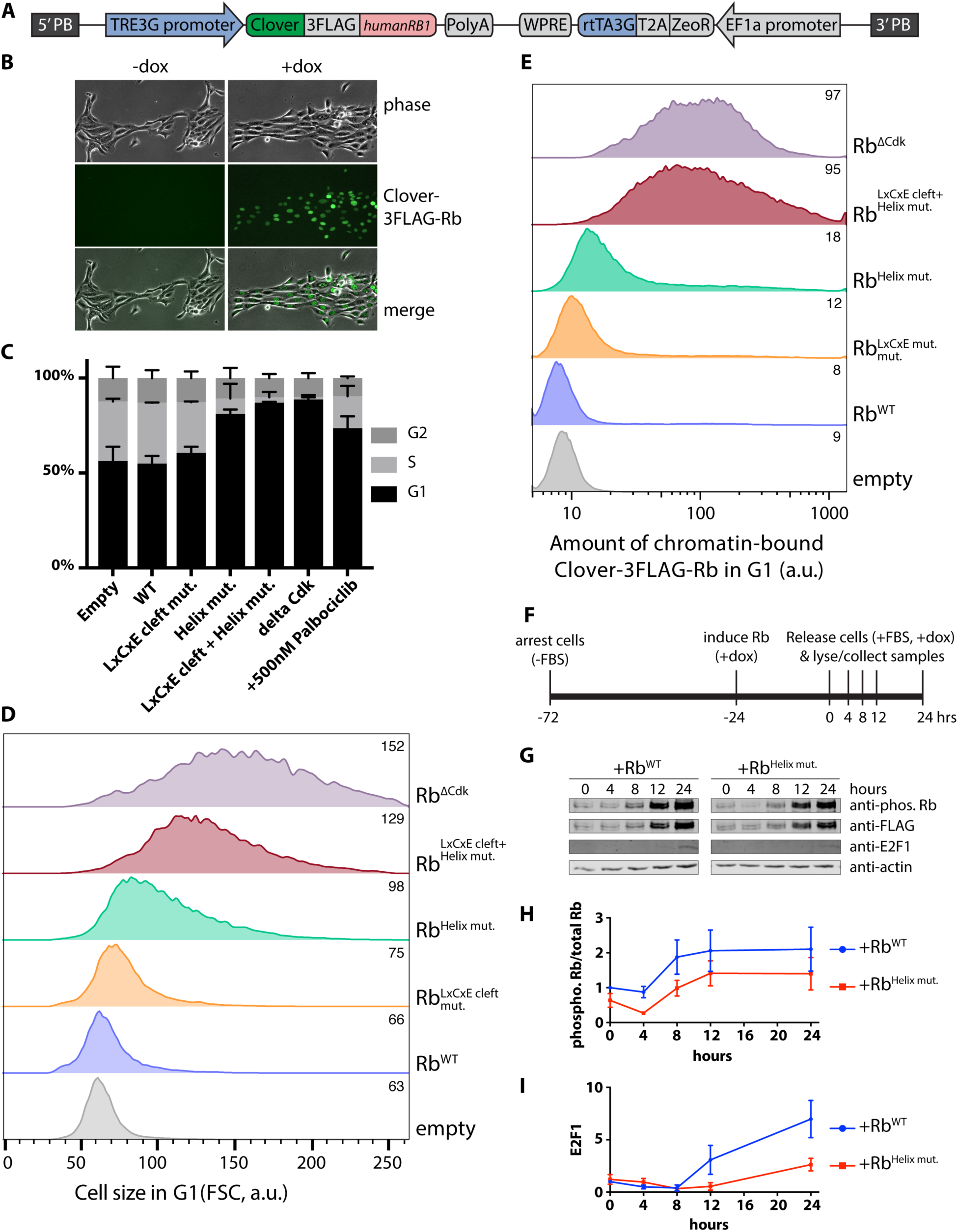
The cyclin D-Rb interaction promotes the G1/S transition, Rb dissociation from chromatin, and E2F1 activation. (A) Map of PiggyBac integration constructs containing a doxycycline-inducible human *RB1* gene fused to fluorescent Clover and 3X FLAG affinity tag sequences. (B) Composite phase and fluorescence images showing expression of Clover-3X FLAG-Rb in HMECs with and without 500 ng/mL doxycycline. (C-E), (C) cell cycle analysis by EdU incorporation and DAPI staining (data are mean ± SEM), (D) cell size in G1, and (E) amount of chromatin-bound Rb in G1 in HMECs expressing Clover-3FLAG alone (empty), wild-type Rb, or the indicated mutant Rb following a 48-hour induction with doxycycline (200 ng/mL). Rb^LxCxE cleft mut.^ lacks the LxCxE docking cleft. Rb^Helix mut.^ denotes an Rb variant where the predicted docking interface residues F897, L901, and R908 are substituted with alanines. Rb^LxCxE cleft+Helix mut.^ contains both indicated mutations. Rb^ΔCdk^ lacks the 15 Cdk phosphorylation sites. Listed in the upper right of each histogram are the medians for (D) and (E). (F) Schematic of Rb phosphorylation timecourse experiment. T98G cells were arrested by serum starvation (- FBS), Rb was induced, and samples were collected at 0, 4, 8, 12, and 24 hours after release (+10% FBS). (G) Immunoblot analysis of lysates from Rb phosphorylation timecourse described in (F) with the denoted antibodies. (H-I) Quantification of (H) phospho-Rb (807/811) over total Clover-3FLAG-Rb and (I) E2F1 from Rb phosphorylation timecourse in (G). Data are mean ± SEM. (C) *n* = 2; (D-E), (G), representative experiments shown (out of three independent experiments); (H-I) *n* = 3.

Next, we examined the phosphorylation of exogenously expressed wild-type Rb and Rb^Helix mut.^ in T98G cells that were arrested and synchronously released into the cell cycle following serum starvation (Figure 5F). Cells expressing Rb^Helix mut.^ displayed less phosphorylated Rb and had weaker E2F1 expression, suggesting that cyclin D-Cdk4,6-dependent phosphorylation during G1 is important for E2F1-dependent transcription and cell cycle entry (Figure 5G-I). We chose to examine E2F1 expression because it is a target of activating E2F transcription factors (Johnson et al., 1994). Consistent with Rb-helix docking being important for activating E2F-dependent gene expression, we observed lower E2F1 protein levels in cells expressing the Rb^Helix mut.^ variant compared to wild type Rb (Figure 5I). Taken together, these experiments are consistent with our biochemical results and show the additive effect of disrupting LxCxE- and helix-based docking mechanisms. Moreover, our analysis strongly supports the role of the cyclin D-Rb interaction as a critical driver of cell cycle entry at the G1/S transition.

### Disruption of the cyclin D-Rb interaction slows tumor growth

We next sought to test our model that cyclin D docks the C-terminal Rb helix to promote cell cycle progression in an animal model. To determine if the Rb variant lacking helix-based docking, Rb^Helix mut.^, is a more potent tumor suppressor than wild-type Rb, we integrated a vector containing doxycycline-inducible Rb alleles into a Kras^+/G12D^;Trp53^-/-^ mouse pancreatic ductal adenocarcinoma cell line (Mazur et al., 2015) (Figure 6A, S6C, and S6D). We then allografted these cell lines into NSG mice by subcutaneous implantation. Tumors were allowed to engraft and grow for five days before we induced expression of the exogenous Rb alleles (Figure 6A). The tumor suppressor function of Rb was enhanced in the Rb^Helix mut.^ variant lacking helix-based docking, and was more enhanced by the removal of the Cdk sites as in Rb^ΔCdk^ (Figure 6B; see Figure S7 for fold changes and p-values for all comparisons). At the end of a biological replicate experiment, tumors were extracted and weighed, further demonstrating the difference between the tumor suppressor potency of wild-type Rb and Rb^Helix mut.^ (Figure 6C). Interestingly, we did not find an additive effect when an LxCxE cleft mutation, Rb^LxCxE cleft mut.^, was combined with either wild-type Rb or Rb^Helix mut.^ (Figure S7), consistent with the previously reported mild phenotype of this mutation (Dick et al., 2000). Our results here support the model in which the cyclin D-Rb helix docking interaction drives Rb inactivation and cell cycle progression *in vivo*.

**Figure 6.**
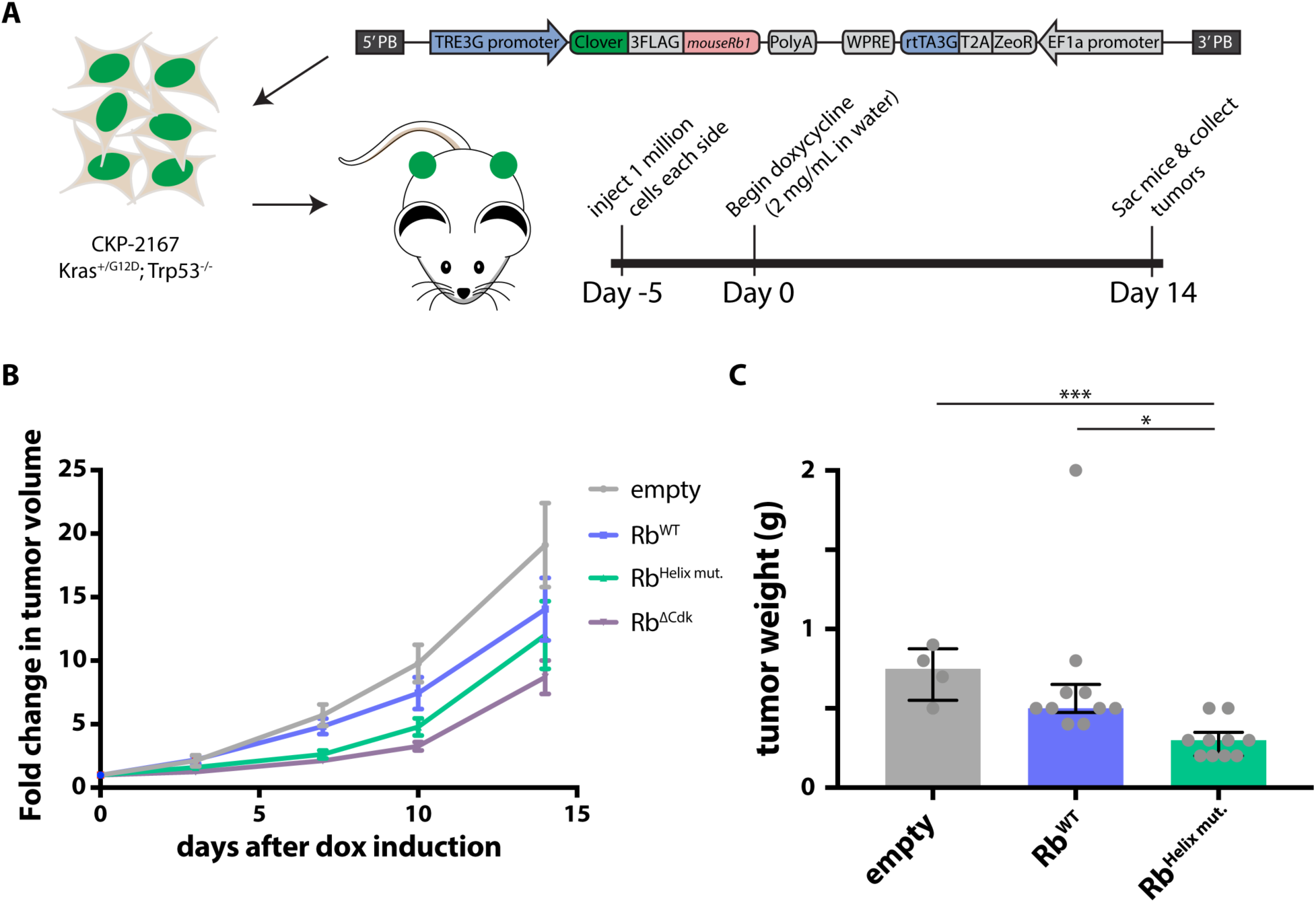
Disruption of the cyclin D-Rb interaction slows tumor growth. (A) Schematic of mouse experiment. PiggyBac integration constructs containing doxycycline-inducible mouse *Rb1* fused to fluorescent Clover and 3FLAG affinity tag sequences were transfected into Kras^+/G12D^; Trp53^-/-^ mouse pancreatic ductal adenocarcinoma cells (CKP-2167). Approximately 1 million CKP-2167 cells expressing variants of doxycycline-inducible mouse Rb were allografted by subcutaneous implantation. After five days of engraftment and growth, mice were given water supplemented with doxycycline (2 mg/mL) for two weeks. (B) Fold change in tumor volume compared to day 0 calculated from caliper measurements. A table of p-values for all fold-change comparisons can be found in Figure S7B. Data shown are mean ± SEM. (C) Median tumor weight 15 days after doxycycline induction of either empty vector, wild-type Rb, or Rb^Helix mut.^. n.s., *, *** denote P > 0.05, P ≤ 0.05, and P ≤ 0.001 respectively. (B) *n* = 12 for empty, Rb^WT^, Rb^Helix mut.^, and Rb^ΔCdk^; (C) *n* = 4 for empty, *n* = 10 for Rb^WT^, and *n* = 10 for Rb^Helix mut.^.

## Discussion

Taken together, our
work demonstrates the importance of helix-based cyclin D docking on Rb to promote phosphorylation, drive cell cycle progression, and inhibit Rb’s tumor suppressive function. In this study, we set out to resolve the conflict between the longstanding model of the G1/S transition and recent reports that cyclin D-Cdk4,6 may drive cell proliferation and survival through Rb-independent mechanisms. We identified a C-terminal helix in Rb that specifically docks cyclin D (Figure 7A). Mutation of the Rb C-terminal helix disrupts its interaction with cyclin D, but maintains the ability of other cyclin-Cdk complexes to phosphorylate Rb. Expressing Rb protein variants lacking the docking helix arrested cells in G1 and slowed tumor growth even though cyclin D-Cdk4,6 could still interact with all of its other targets. A model where cyclin D inhibits Rb indirectly through downstream cyclins is therefore unlikely because these Rb variants lacking the docking helix are readily targeted by downstream cyclins E and A. Thus, Rb phosphorylation by cyclin D-Cdk4,6 is a crucial first step in driving the G1/S transition.

**Figure 7.**
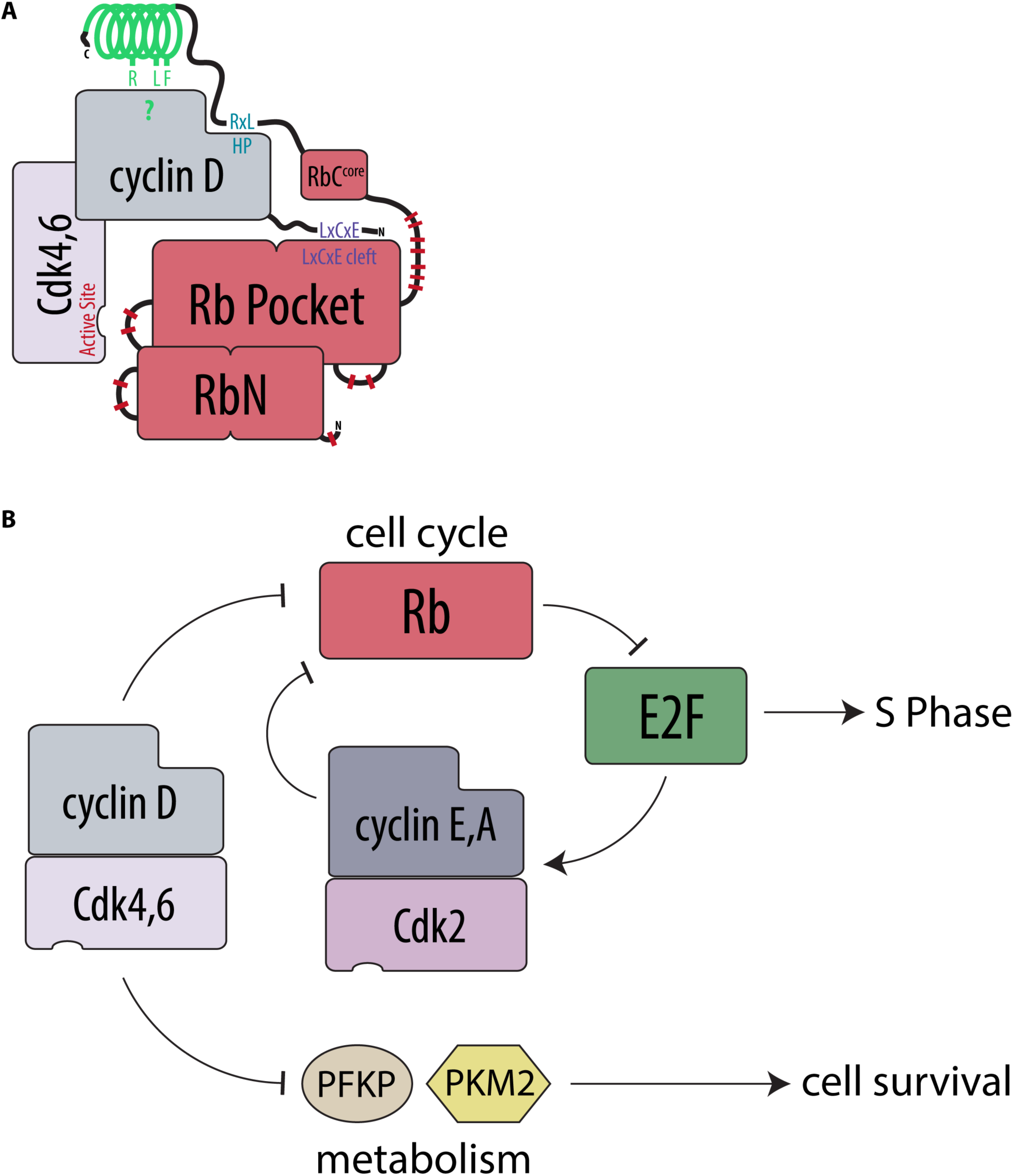
Cyclin D-Cdk4,6 inactivates Rb through multiple, specific docking interactions to drive cell cycle progression. (A) Schematic of the multiple docking interactions between cyclin D and Rb. (B) Model of the major functions of cyclin D-Cdk4,6 complexes.

While Rb is clearly an important target for cyclin D to promote cell proliferation, it may not be the only such target. While we believe the major cell cycle function of cyclin D-Cdk4,6 is to inactivate and phosphorylate Rb through helix-based docking, the complex also independently influences cell survival by phosphorylating and inactivating metabolic enzymes (Figure 7B). Thus, cell proliferation *in vivo* may be enhanced by increasing cell survival *and* by driving cell division through distinct cyclin D-Cdk4,6 targets. By examining Rb-docking helices across metazoans, we identified a consensus helix sequence motif, which we then used to generate a list of potential substrates in the human proteome (Table S1). This list may provide more insight into non-canonical substrates and functions of cyclin D-Cdk4,6 that could promote proliferation.

In general, progression through the eukaryotic cell cycle is characterized by a monotonic increase in cyclin-dependent kinase activity (Stern and Nurse, 1996). However, not all cell cycle-dependent substrates are phosphorylated at the same time (Swaffer et al., 2016; 2018). Early substrates are typically targets of early cyclin-Cdk complexes via specific docking mechanisms (Loog and Morgan, 2005). To prevent later substrates from being phosphorylated by the early-activated cyclin-Cdk complexes, these early complexes are typically characterized by lower intrinsic kinase activity. For example, in budding yeast, peptides containing a single Cdk site derived from histone H1 were phosphorylated at progressively higher rates by G1-phase, S-phase, and mitotic cyclins (Kõivomägi et al., 2011). Here, we report a similar progressive increase in intrinsic kinase activity towards histone H1 by animal cyclin-Cdk complexes. By far, the lowest activity towards H1 was found in cyclin D-Cdk4,6 complexes. Thus, the progression of the cell cycle from high-specificity cyclin-Cdk complexes through to high-activity complexes is likely a conserved feature of eukaryotic cell cycle control.

The low activity of cyclin D-Cdk4,6 complexes may be compensated for by its specific docking mechanism. In contrast to previously identified cyclin-Cdk docking motifs, which are short linear motifs in intrinsically disordered regions of proteins, Rb helix-based docking is unique in its structural requirement. While short linear motifs are easy to evolve (Davey et al., 2015), the addition of a structural requirement, such as a helix, likely makes it much more difficult. This is because both the helix and the appropriate interface residues must align. The difficulty of evolving helix-based docking could therefore underlie the observation that cyclin D-Cdk4,6 appears to have relatively few substrates compared to other cyclin-Cdk complexes (Malumbres and Barbacid, 2005). We speculate that cyclin D-Cdk4,6 has few substrates so that this low-activity kinase can be able to adequately phosphorylate its key targets, including Rb.

Cyclin D-Cdk4,6 complexes clearly play a role in cancer, and a series of cancer drugs targeting the ATP pockets of Cdk4 and Cdk6 are successfully emerging from clinical trials (Sherr et al., 2016). However, while Cdk4,6 inhibitors have been observed to limit disease progression, overall survival is not affected for drugs such as palbociclib at the current follow-up times (Finn et al., 2016). That Cdk4,6 inhibitors have significant off-target activities (Hafner et al., 2017) raises the possibility that cancer therapies can be improved by a new class of drugs targeting cyclin substrate recognition based on our helix docking mechanism.

## Acknowledgments

We thank Nick Dyson, Ioannis Sanidas, and Joe Lipsick for their thoughtful comments on the manuscript, Mart Loog for discussions on cyclin docking, and Nick Buchler for discussions on Rb evolution. We thank Ali Shariati for dox-inducible PiggyBac reagents, Steve Elledge for HMEC-hTERT1 cells, Robert Weinberg and Steve Dowdy for Rb plasmids, Martha Cyert for advice and reagents for the GST binding assay, and Monte Winslow for NSG mice. Flow cytometry data were collected on an instrument in the Stanford Shared FACS Facility obtained using the NIH S10 Shared Instrument Grant S10RR027431-01. This work was supported by the NIGMS through R01 GM092925 and R01 GM115479, and the NCI through the Cancer Biology Training Grant T32 CA009302.

## Author contributions

B.R.T., E.Z., S.M.R., J.S., M.K., and J.M.S. conceived and designed the experiments. B.R.T., E.Z., S.C., C.S.T., and M.K. performed the experiments. S.X. performed the computational analysis. B.R.T., M.K., and J.M.S. wrote the manuscript.

## Materials and Methods

### Protein expression and purification

Full-length, N-terminally glutathione S-transferase-tagged (GST-tagged), Rb family proteins were expressed in the *E. coli* strain BL21, purified by glutathione-agarose affinity chromatography (Sigma-Aldrich G4510), and eluted with 50mM Tris pH 8.0; 100mM KOAc; 25mM MgOAc; 10% glycerol; 15mM Glutathione. Histone H1 protein used as a general substrate for Cdk was purchased from EMD Millipore (14-155).

GST-Cdk phosphorylation site fusion proteins (Table S1 and Figure S4) containing a GST-tag, a TEV-protease cleavage site, and a single Cdk phosphorylation site peptide were expressed and purified as described above. The three different Cdk phosphorylation sites that we used were (1) a Cdk site patterned after a region of similar sequence found in histone H1 protein (Hagopian et al., 2001; Kõivomägi et al., 2011) (H1 site: PKTPKKAKKL), (2) an S780 Cdk site from Rb(Kitagawa et al., 1996) (Rb 775-787: RPPTLSPIPHIPR), and (3) an S795 Cdk site from Rb (Grafström et al., 1999) (Rb 790-805: YKFPSSPLRIPGGNIY). To test docking specificity of the cyclin-Cdk complexes, we also fused these GST-Cdk site fusion proteins to the Rb C-terminal Helix (Rb 895-915: SKFQQKLAEMTSTRTRMQKQK) or the Cdc6 RxL docking motif (Takeda et al., 2001) (Cdc6 89-103: HTLKGRRLVFDNQLT) using a G_4_S glycine-serine linker (GGGGS).

Human cyclin-Cdk fusion complexes were purified from budding yeast cells (Schwarz et al., 2018) using a 3X FLAG affinity purification method, modified from a previous protocol used for HA-tag purification (McCusker et al., 2007). Briefly, N-terminally tagged cyclin-Cdk fusions were cloned into 2-micron vectors using a glycine-serine linker (Rao et al., 1999) and overexpressed from the *GAL1* budding yeast promoter. The overexpressed 3FLAG-tagged cyclin-Cdk complexes were then purified by immunoaffinity chromatography using ANTI-FLAG M2 affinity agarose beads (Sigma-Aldrich A2220) and eluted with 0.2 mg/mL 3X FLAG peptide (Sigma-Aldrich F4799). We note that similar cyclin-Cdk fusions have previously been able to restore wild-type function *in vivo* (Chytil et al., 2004). Plasmids and proteins used in this study are listed in Table S3. Yeast strains are listed in Table S4.

### In vitro kinase assays

For all experiments, equal amounts of substrate and purified kinase complexes were used. Substrate concentrations were kept in the range of 1-5 µM for different experiments, but did not vary within any experiment. Reaction aliquots were taken at two time points (8 and 16 minutes) and the reaction was stopped with SDS-PAGE sample buffer. The basal composition of the assay mixture contained 50 mM HEPES pH 7.4, 20 mM Tris pH 8.0, 150 mM NaCl, 5 mM MgCl_2_, 10 mM MgOAc, 40 mM KOAc, 6 mM glutathione, 0.2 mg/ml 3X FLAG peptide, 6% glycerol, 3 mM EGTA, 0.2 mg/ml BSA and 500 µM ATP (with 2 µCi of [γ-^32^P] ATP added per reaction; PerkinElmer BLU502Z250UC). Phosphorylated proteins were separated on 10% SDS-PAGE gels. Phosphorylation of substrate proteins was visualized using autoradiography (Typhoon 9210; GE Healthcare Life Sciences). Autoradiographs were quantified with the ImageQuant TL Software. We note that phosphorylation of H1 and Rb by our cyclin-Cdk fusions occurs in the linear range for the 8 and 16 minute time-points (Figure S1A-H).

### GST Binding assay

GST-tagged Rb proteins were dialyzed to remove glutathione using a buffer containing 50mM Tris pH 8.0, 100mM KOAc, 25mM MgOAc, and 10% glycerol. Next, GST-Rb proteins were bound to glutathione-agarose beads (Sigma-Aldrich G4510) for 1 hour at 4°C in a binding buffer containing 50 mM Tris pH 8.0, 150 mM NaCl, 1% triton X-100, and 1 mM DTT. This bead-GST-Rb mixture was washed with this binding buffer and incubated with an equimolar amount of 3FLAG-tagged cyclin-Cdk complexes for 2-3 hours at 4°C. Beads were then washed with binding buffer and eluted with 2X SDS-PAGE sample buffer. Input and pulldown samples were then analyzed by immunoblotting.

### Circular Dichroism

Circular dichroism (CD) spectra were recorded using a Jasco J-1500 CD spectrometer. Samples contained 30 µM recombinant Rb 890-920 in a buffer containing 25 mM sodium phosphate and 100 mM NaCl (pH 6.1). The data were fit by calculating a weighted average of reference spectra measured for poly-L-lysine, which adopts known different secondary structures depending on pH and temperature (Greenfield and Fasman, 1969).

### Helix prediction, bioinformatics analysis, and helix motif search

Secondary structure predictions were carried out on the PSIPRED protein structure prediction server using the PSIPRED v3.3 Secondary Structure prediction method (Buchan et al., 2013) (http://bioinf.cs.ucl.ac.uk/psipred/). Helical wheel projections of predicted C-terminal helices were generated using the “Helical Wheel Projections” tool (http://rzlab.ucr.edu/scripts/wheel/wheel.cgi).

The full-length sequences of Rb family members (Medina et al., 2016) were aligned by MAFFT-L-INS-I (Katoh et al., 2017). A profile Hidden-Markov model (HMM) was generated using the HMMER3 web service (Finn et al., 2015). JackHMMER used to iteratively search for Rb homologs based on the profile HMM, using a stringent E-value cutoff of 1e-10 against UniProt Reference Proteomes. Of the total 1072 eukaryotic Rb homologs retrieved, 682 were metazoan. All sequences were re-aligned using MAFFT-L-INS-i (maxitr=1000). The aligned metazoan sequences were then trimmed to focus on sequence positions occupied by human Rb. Membership in *RB1*, *RBL1*, or *RBL2* sub-families were determined by manual examination of the phylogenetic tree generated using FastTree (Price et al., 2009). Non-metazoan Rb sequence from taxa closely related to metazoan were examined manually.

Aligned sequences from the helix region of metazoan RB family members were used to generate a position specific weight matrix (PSSM). The PSSM was used to score protein sequences in the UniProtKB/Swiss-Prot reviewed human proteome (UP000005640). PSIPRED was used perform helicity prediction on sequences with PSSM scores above 20, which corresponds to 5 standard deviation above the mean random score distribution). The sequences were further filtered by the presence of CDK substrate sites ([S/T]P) within the full-length protein. All analysis was done using BioPython (Cock et al., 2009). Results from helix motif search are listed in Table S2.

### Doxycycline-inducible PiggyBac integration vector construction

We cloned an Rb expression cassette, driven by the *TRE3G* doxycycline-inducible promoter (Clontech 631168), into a PiggyBac integration plasmid containing 5’ and 3’ PiggyBac homology arms (Ding et al., 2005; Shariati et al., 2018). The expression cassette contains the human *RB1* or mouse *Rb1* genes fused to fluorescent Clover and 3FLAG affinity tag sequences, a zeocin resistance gene, and a Tet-On 3G transactivator gene driven by the *Ef1α* promoter. Plasmids are listed in Table S3.

### Cell culture and cell line construction

Immortalized human mammary epithelial cells (HMEC-hTERT1 cells, also referred to as HMECs) from the Elledge lab (Sack et al., 2018) were cultured in MEGM™ Mammary Epithelial Cell Growth Medium (Lonza CC-3150). T98G and Kras^+/G12D^; Trp53^-/-^ mouse pancreatic ductal adenocarcinoma cells (CKP-2167) were cultured in DMEM supplemented with 10% FBS, 4.5g/L Glucose, 2 mM L-glutamine, and Sodium Pyruvate (Mazur et al., 2015). Cell lines stably expressing doxycycline-inducible Rb variants were generated by transfecting cells plated into individual wells of a 6 well dish with 2.2 µg of doxycycline-inducible PiggyBac integration plasmid and 1.1 µg of PiggyBac Transposase plasmid (Ding et al., 2005) using the FuGene HD reagent (Promega E2311). Zeocin (400ug/mL) selection began two days after transfection. Zeocin resistant cells were maintained as polyclonal cell lines. Prior to flow cytometry and immunoblot analysis, cells were grown in the presence of doxycycline (500 ng/mL) for two days. Cell lines are listed in Table S4.

### Flow cytometry

Flow cytometry analysis was performed on BD LSRII.UV cytometer. Live cells were prepared for flow cytometry by washing with 1X PBS, trypsinizing, and resuspending in 1X PBS. For live cells, we measured forward scatter area as a readout for cell size. For measurement of newly synthesized DNA by nucleoside analog incorporation, cells were incubated with 10 µM EdU for 30 minutes at 37°C, fixed in 3% formaldehyde for 10 minutes at 37°C, and permeabilized with 90% ethanol for 30 minutes on ice. EdU was detected using the Click-iT™ Plus EdU Alexa Fluor™ 594 Flow Cytometry Assay Kit (C10646) and cells were stained with 3 µM DAPI for 10 minutes at room temperature. For these fixed cells, we measured EdU, DAPI fluorescence as a readout for DNA content, Clover green protein fluorescence as a readout for the amount of tagged exogenous Rb, and forward scatter area as a readout for cell size. To measure the amount of DNA-associated Rb, we used a soluble protein extraction method (Håland et al., 2015). Briefly, the cells were harvested by trypsinization and pelleted by centrifugation. The cell pellet was then resuspended in ice-cold low salt extraction buffer (0.1% Igepal CA-630, 10 mM NaCl, 5 mM MgCl2, 0.1 mM PMSF, 10 mM Potassium phosphate buffer pH 7.4) and incubated on ice for 1 minute. Then, the cells were fixed by adding paraformaldehyde to a final concentration of 3% and incubating on ice for 1 hour. Fixed cells were washed once with 1X PBS and then stained with 20 µM Hoechst 33342 DNA dye for 10 minutes at 37°C in 1X PBS. DNA content and DNA-bound Clover-tagged Rb amounts were measured with a BD LSRII.UV flow cytometer.

### Immunoblotting

Portions of harvested tumors were resuspended at 100 mg/mL in RIPA buffer supplemented with protease and phosphatase inhibitors. Next, we homogenized these tumor samples with pestles in tubes, and then sonicated the samples for 10 seconds at 50% amplitude on a Fisherbrand Model 120 Sonic Dismembrator. Cells cultured on dishes were collected by scraping in 1X PBS and lysed in RIPA buffer supplemented with protease and phosphatase inhibitors. Proteins from lysates were separated on a 10% SDS-PAGE gel and transferred to a nitrocellulose membrane using the iBlot 2 dry blotting system (Invitrogen IB21001).

Membranes were incubated overnight at 4°C with the following antibodies: Phospho-Rb (Ser807/811) (D20B12) XP^®^ Rabbit mAb (Cell Signaling Technology #8516), Purified Mouse Anti-Human Retinoblastoma Protein Clone G3-245 (BD Biosciences 554136), Monoclonal ANTI-FLAG^®^ M2 antibody produced in mouse (Sigma-Aldrich F1804), E2F-1 Antibody (Cell Signaling Technology #3742), and mouse monoclonal β-Actin Antibody N-21 (Santa Cruz Biotechnology sc-130656). The primary antibodies were detected using the fluorescently labeled secondary antibodies IRDye^®^ 680LT Goat anti-Mouse IgG (LI-COR 926-68020) and IRDye^®^ 800CW Goat anti-Rabbit IgG (LI-COR 926-32211). Membranes were imaged on a LI-COR Odyssey CLx and analyzed with LI-COR ImageStudio software.

### Subcutaneous tumor implantation

CKP-2167 expressing variants of doxycycline-inducible mouse Rb were allografted by subcutaneous implantation in NSG mice. For each implantation, approximately 1 million cells were suspended in 100 µL of 1X PBS and mixed with 100 µL of Matrigel^®^ Basement Membrane Matrix (Corning 356237). This mixture was injected into the left and right flanks of each mouse. After five days of engraftment and growth, mice were given water supplemented with doxycycline (2 mg/mL) for two weeks. Tumor lengths and widths were measured with calipers and volumes were estimated with the formula V = (4/3) × π × r^3^, where r is half of the average tumor diameter or (length + width) ÷ 4. At the end of the experiment, mice were sacrificed, and tumors were harvested.

## Supplemental Figures

**Figure S1.**
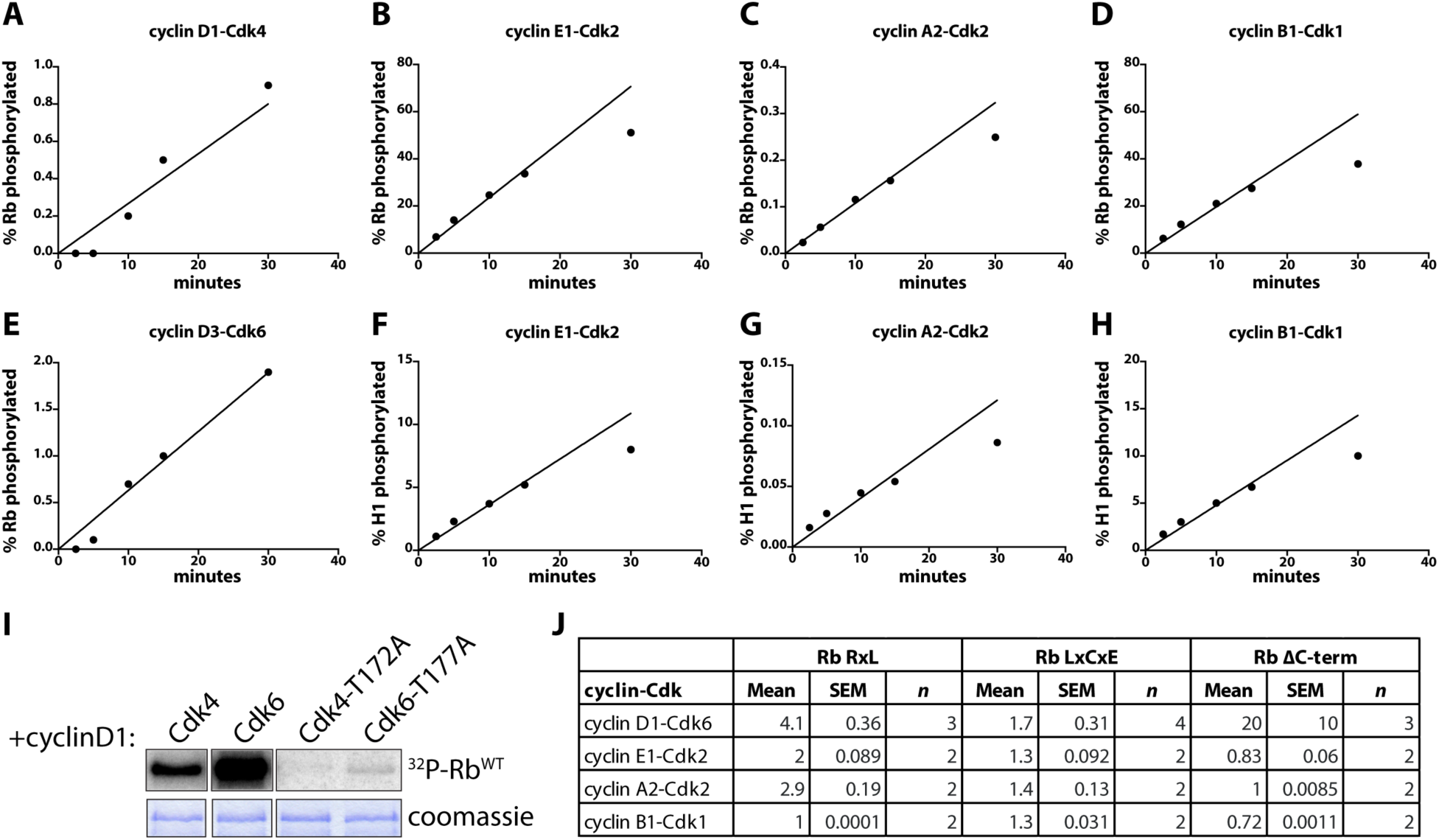
Characterization of cyclin-Cdk fusion kinases. (A-H) *In vitro* kinase assay timecourses of the denoted cyclin-Cdk complexes with Rb or histone H1 as a substrate, as indicated on the y-axis. Kinase assay samples were taken at 2.5, 5, 10, 15, and 30 minutes. To calculate percentage of substrate phosphorylated for a given timepoint, Rb and H1 were fully phosphorylated by adding maximum amounts of cyclin A2-Cdk2 and cyclin B1-Cdk2, respectively. Each timepoint was quantified divided by the maximum substrate phosphorylation. Lines were fit to the first four timepoints and forced through the origin. We note that these kinase reactions stay within the linear range for the timepoints used in all other experiments. (I) *In vitro* kinase assays comparing phosphorylation of Rb by wild-type kinases (cyclin D1-Cdk4 and cyclin D1-Cdk6) and T-loop mutant kinases (cyclin D1-Cdk4-T172A and cyclin D1-Cdk6-T177A). The coomassie stained gels showing equal amounts of substrate used in each reaction are placed below each autoradiograph. Fold change of WT to T-loop mutant is 230±20 for Cdk4 and 210±30 for Cdk6. (J) Mean fold changes, SEM, and *n*, for the denoted Rb variant phosphorylation by the denoted cyclin-Cdk complexes. (A-H), representative experiments shown (out of two independent experiments); (I), *n* = 2.

**Figure S2.**
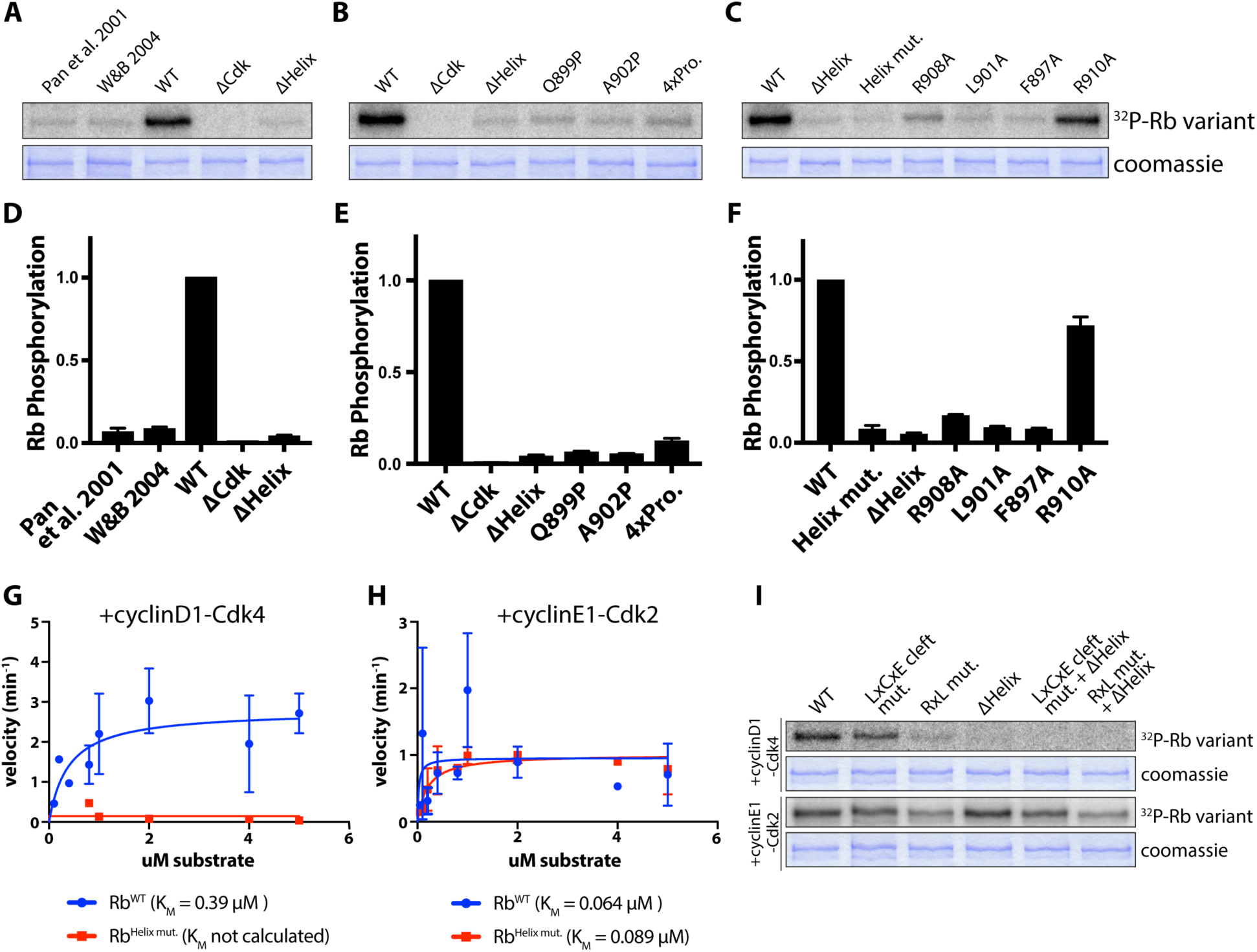
Cyclin D-Cdk4,6 complexes dock the Rb C-terminal helix. (A-C) *In vitro* kinase assays of the indicated Rb variants. “Pan et al. 2001” denotes an Rb-L901Q variant and “W&B 2004,” reported in Wallace & Ball 2004, denotes an Rb-F897A-Q899A-M904A-R908A-R910A variant. ΔCdk denotes an Rb variant lacking the 15 Cdk phosphorylation sites. ΔHelix denotes an Rb variant that lacks the C-terminal helix amino acids 895-915. 4xPro. denotes an Rb variant with proline substitutions at the K896, Q898, Q899, and A902 residues within the C-terminal helix. Helix mut. denotes an Rb variant where the predicted docking interface residues F897, L901, and R908 are substituted with alanines. The coomassie stained gel showing equal amounts of substrate used in each reaction is placed below each autoradiograph. (D-F) Quantification of the *in vitro* kinase assays shown in (A-C), respectively. Data are mean ± SEM. (G and H) Michaelis-Menten K_M_ calculation for (G) cyclin D1-Cdk4 or (H) cyclin E1-Cdk2 with Rb^WT^ or Rb^Helix mut.^. *In vitro* kinase assays were performed with a range of Rb substrate concentrations from 0.05 µM to 5 µM and samples were taken at 8 and 16 minutes. The Michaelis-Menten K_M_ constants were calculated in GraphPad Prism 7. (I) *In vitro* kinase assays of the denoted Rb variant with cyclin D1-Cdk4 or cyclin E1-Cdk2. LxCxE cleft mut. lacks the LxCxE docking cleft. RxL mut. lacks all C-terminal RxL sequences that dock cyclin hydrophobic patches. The coomassie stained gels showing equal amounts of substrate used in each reaction is shown below each autoradiograph. (L) cyclin E1-Cdk2. (A-H), *n* = 2; (I) representative experiments shown (out of two independent experiments).

**Figure S3.**
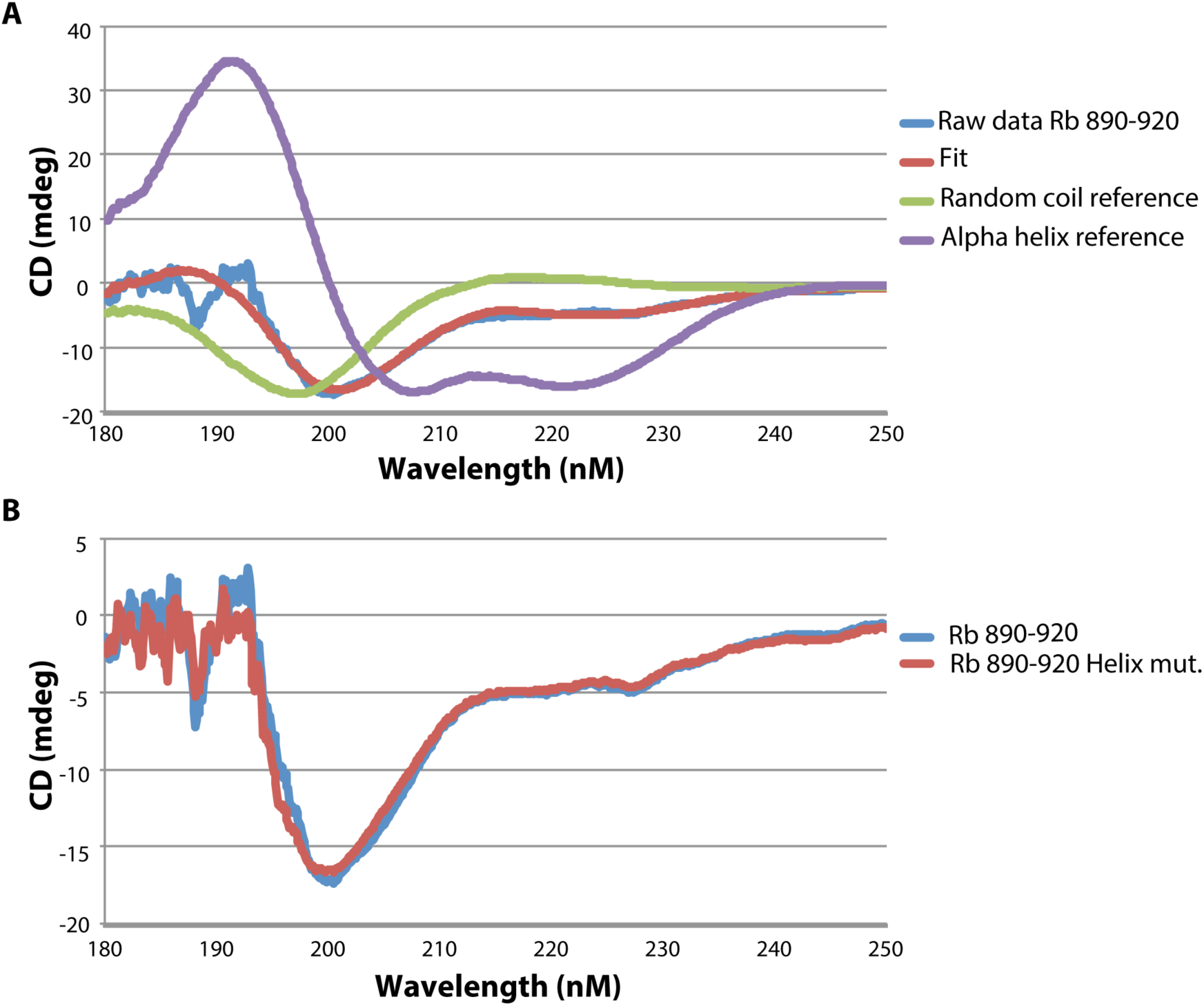
Circular dichroism supporting helical propensity in Rb 890-920. (A) Circular dichroism (CD) spectrum of recombinant 30 µM Rb 890-920 in a buffer containing 25 mM sodium phosphate and 100 mM NaCl (pH 6.1). Spectra were recorded using a Jasco J-1500 CD Spectrometer. The data were fit by calculating a weighted average of reference spectra measured for poly-L-lysine. The best fit of the data indicates that the peptide contains 65% random coil and 35% alpha helix. The reference spectra for helix and random coil are shown for comparison. Considering our secondary structure prediction, we suggest that the peptide alone in solution exists in a conformational equilibrium with some tendency to form helical structure. (B) Comparison of CD spectra for wild type (30 µM) and Helix mut. (20 µM) Rb 890-920. Helix mut. denotes an Rb variant where the predicted docking interface residues F897, L901, and R908 are substituted with alanines.

**Figure S4.**
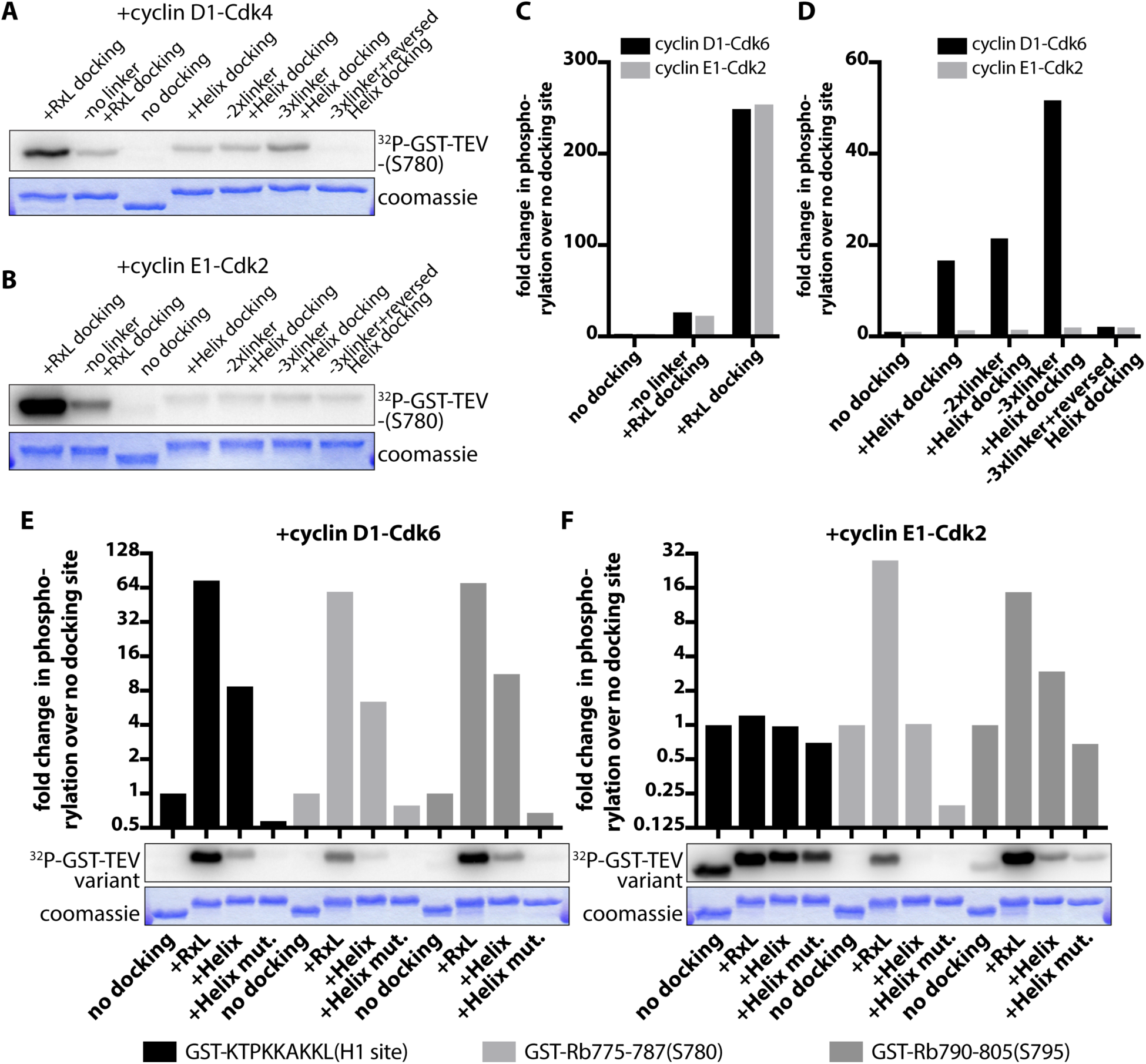
Characterization of engineered GST-Cdk phosphorylation site fusion proteins. (A-D) *In vitro* kinase assays of the denoted engineered GST-Cdk phosphorylation site fusion protein with (A) cyclin D1-Cdk6 or (B) cyclin E1-Cdk2. Full sequences can be found in Table S1. The coomassie stained gels showing equal amounts of substrate used in each reaction are placed below each autoradiograph. The *in vitro* kinase assays in (A and B) were quantified with respect to (C) RxL docking or (D) Helix docking. (E-F) *In vitro* kinase assays of the denoted engineered GST-Cdk phosphorylation site fusion protein with different phosphorylation and docking sites using (E) cyclin D1-Cdk6 or (F) cyclin E1-Cdk2. The coomassie stained gels showing equal amounts of substrate used in each reaction are placed below each autoradiograph. (A-F), *n* = 1.

**Figure S5.**
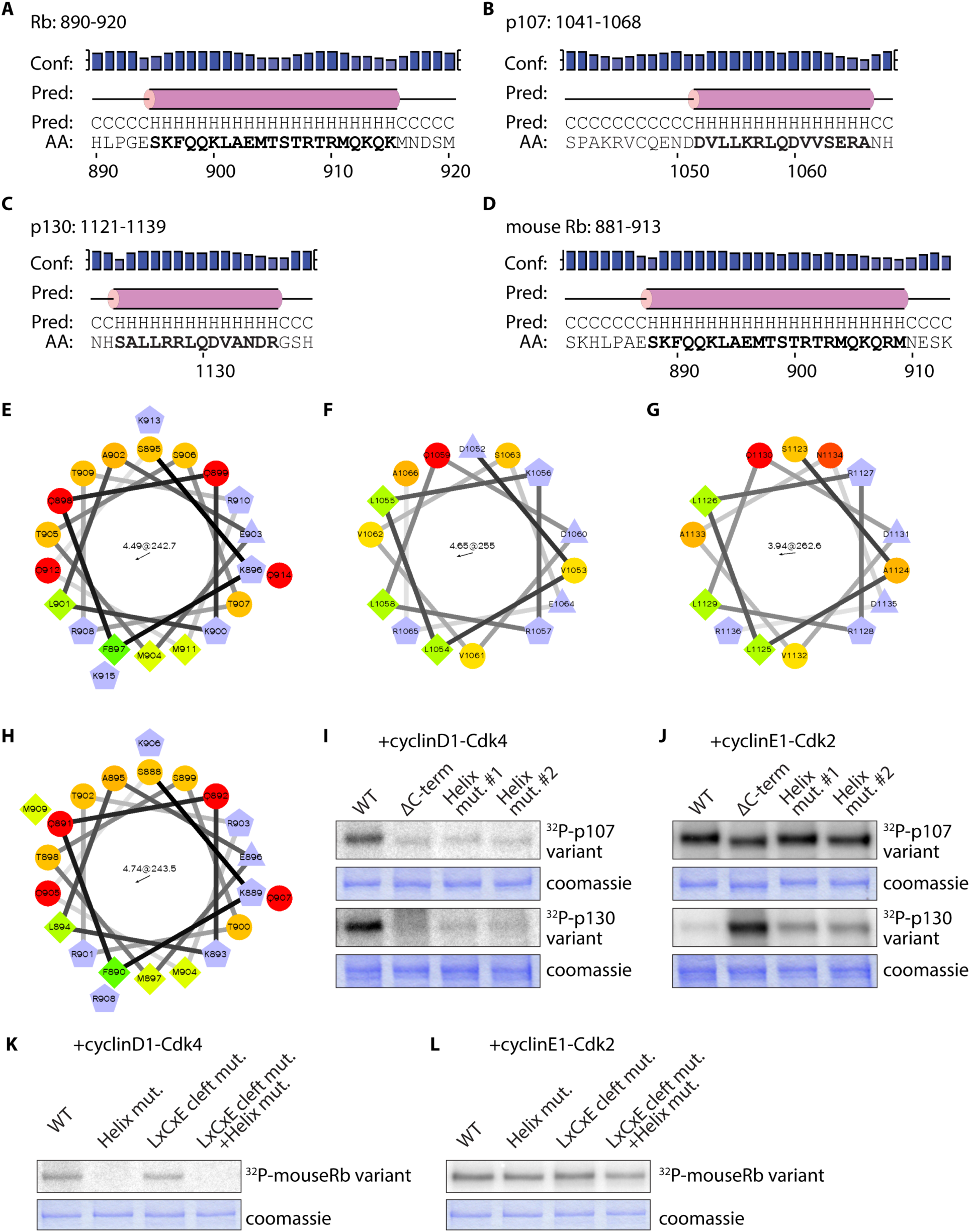
C-terminal helices in the Rb family proteins. (A-D) Secondary structure predictions were generated for the C-termini of (A) human Rb, (B) human p107, (C) human p130, and (D) mouse Rb. (E-H) Helical wheel projections, based on the sequence predictions in (A-D), were generated for the C-terminal helices of (E) human Rb, (F) human p107, (G) human p130, and (H) mouse Rb. (I and J) *In vitro* kinase assays of the denoted p107 or p130 variant with (I) cyclinD1-Cdk4 or (J) cyclinE1-Cdk2. ΔC-term denotes truncating p107 at amino acid position 1014 and p130 at amino acid position 1122. Helix mut. #1 denotes alanine substitution of L1054, L1058, and R1065 of p107 and L1125, L1129, and R1136 of p130. Helix mut. #2 denotes alanine substitution of L1058 and R1065 of p107 and L1129 and R1136 of p130. The coomassie stained gels showing equal amounts of substrate used in each reaction are placed below each autoradiograph. (K and L) *In vitro* kinase assays of the denoted mouse Rb variant with (K) cyclinD1-Cdk4 or (L) cyclinE1-Cdk2. Helix mut. denotes alanine substitution of F890, L894, and R901. LxCxE cleft mut. lacks the LxCxE docking cleft. The coomassie stained gels showing equal amounts of substrate used in each reaction are placed below each autoradiograph. (I-L), representative experiments shown (out of two independent experiments).

**Figure S6.**
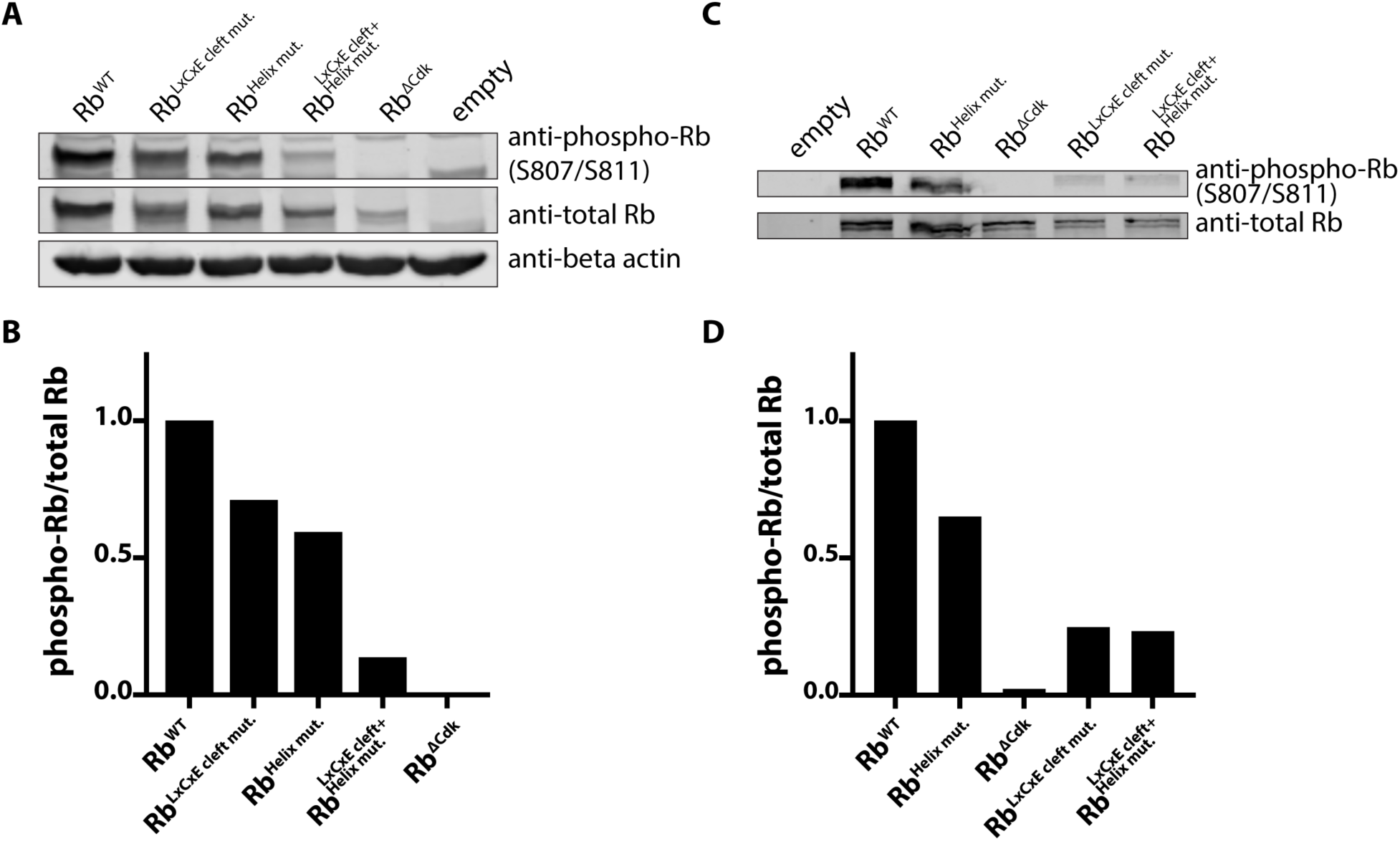
Immunoblot analysis of Rb induction in cell lines. (A) Immunoblot analysis of lysates from HMECs expressing the denoted Rb variants. (B) Quantification of phospho-Rb to total Rb in HMECs. (C) Immunoblot analysis of lysates from CKP-2167 cells expressing the denoted Rb variants. (D) Quantification of phospho-Rb to total Rb in CKP-2167 cells. (A-D), representative experiments shown (out of three independent experiments).

**Figure S7.**
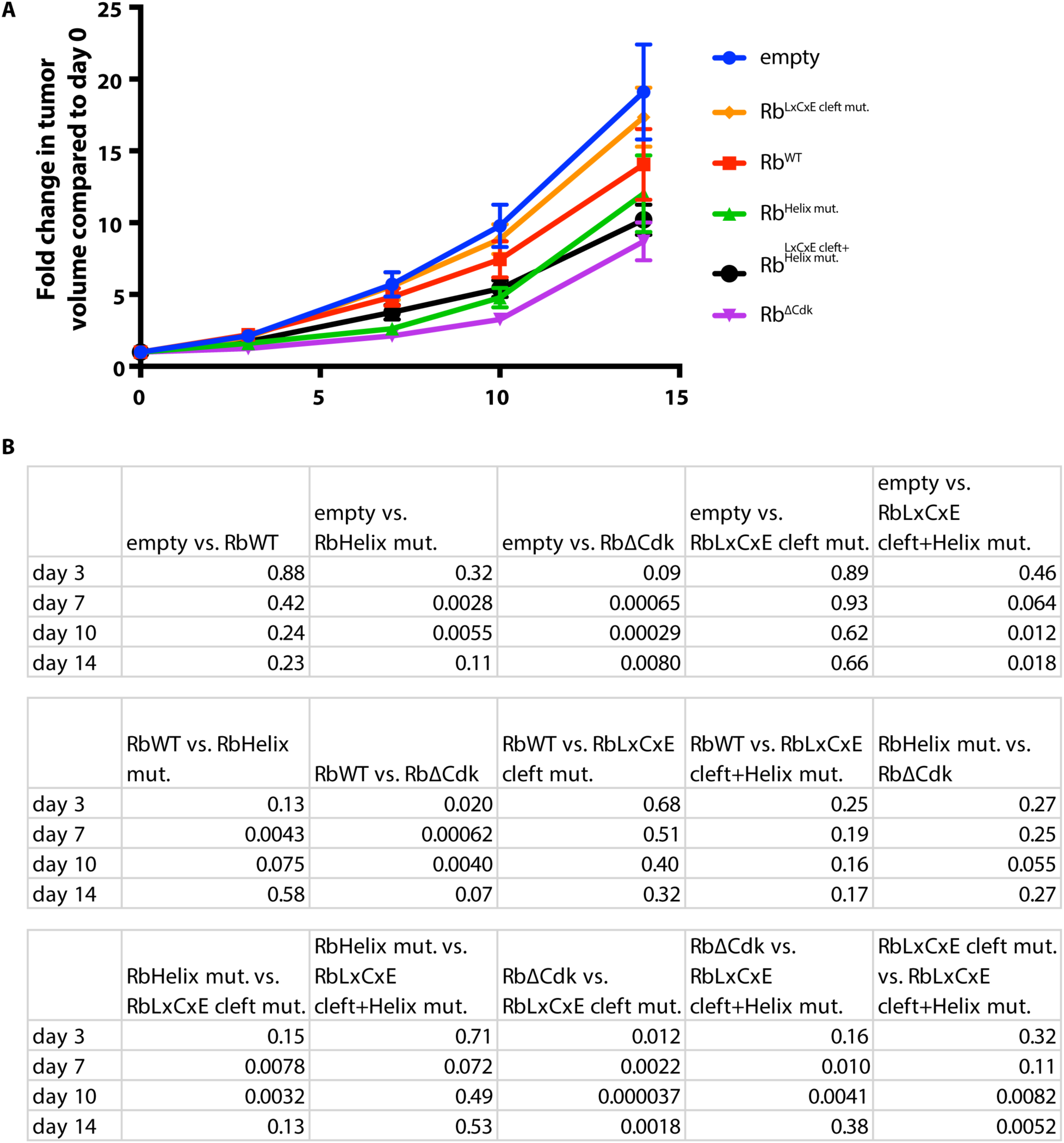
Disruption of the cyclin D-Cdk4,6-Rb interaction slows tumor growth. (A) Fold change in tumor volume compared to day 0 calculated from caliper measurements. Empty denotes a cell line expressing only Clover-3X FLAG. Rb^LxCxE cleft mut.^ denotes an Rb variant that lacks the LxCxE docking cleft. Rb^Helix mut.^ denotes an Rb variant where the predicted docking interface residues F897, L901, and R908 are substituted with alanines. Rb^LxCxE cleft+Helix mut.^contains both indicated mutations. Rb^ΔCdk^ denotes an Rb variant that lacks the 15 Cdk phosphorylation sites. (B) P-values for every fold change in tumor volume comparison. (A) *n* = 12 for each condition.

## References

Adams, P.D., Li, X., Sellers, W.R., Baker, K.B., Leng, X., Harper, J.W., Taya, Y., and Kaelin, W.G. (1999). Retinoblastoma protein contains a C-terminal motif that targets it for phosphorylation by cyclin-cdk complexes. Mol. Cell. Biol. 19, 1068–1080.

Anders, L., Ke, N., Hydbring, P., Choi, Y.J., Widlund, H.R., Chick, J.M., Zhai, H., Vidal, M., Gygi, S.P., Braun, P., et al. (2011). A Systematic Screen for CDK4/6 Substrates Links FOXM1 Phosphorylation to Senescence Suppression in Cancer Cells. Cancer Cell 20, 620–634.

Bertoli, C., Skotheim, J.M., and de Bruin, R.A.M. (2013). Control of cell cycle transcription during G1 and S phases. Nature Publishing Group 14, 518–528.

Bhaduri, S., and Pryciak, P.M. (2011). Cyclin-specific docking motifs promote phosphorylation of yeast signaling proteins by G1/S Cdk complexes. Curr. Biol. 21, 1615–1623.

Bloom, J., and Cross, F.R. (2007). Multiple levels of cyclin specificity in cell-cycle control. Nat Rev Mol Cell Biol 8, 149–160.

Burkhart, D.L., and Sage, J. (2008). Cellular mechanisms of tumour suppression by the retinoblastoma gene. Nat Rev Cancer 8, 671–682.

Cappell, S.D., Chung, M., Jaimovich, A., Spencer, S.L., and Meyer, T. (2016). Irreversible APC(Cdh1) Inactivation Underlies the Point of No Return for Cell-Cycle Entry. Cell 166, 167–180.

Cappell, S.D., Mark, K.G., Garbett, D., Pack, L.R., Rape, M., and Meyer, T. (2018). EMI1 switches from being a substrate to an inhibitor of APC/CCDH1 to start the cell cycle. Nature 558, 313–317.

Cross, F.R., and Jacobson, M.D. (2000). Conservation and function of a potential substrate-binding domain in the yeast Clb5 B-type cyclin. Mol. Cell. Biol. 20, 4782–4790.

Davey, N.E., Cyert, M.S., and Moses, A.M. (2015). Short linear motifs - ex nihilo evolution of protein regulation. Cell Commun. Signal 13, 43.

Di Fiore, B., Davey, N.E., Hagting, A., Izawa, D., Mansfeld, J., Gibson, T.J., and Pines, J. (2015). The ABBA Motif Binds APC/C Activators and Is Shared by APC/C Substrates and Regulators. Developmental Cell 32, 358–372.

Dick, F.A., and Rubin, S.M. (2013). Molecular mechanisms underlying RB protein function. Nature Publishing Group 14, 297–306.

Dick, F.A., Sailhamer, E., and Dyson, N.J. (2000). Mutagenesis of the pRB Pocket Reveals that Cell Cycle Arrest Functions Are Separable from Binding to Viral Oncoproteins. Mol. Cell. Biol. 20, 3715–3727.

Dowdy, S.F., Hinds, P.W., Louie, K., Reed, S.I., Arnold, A., and Weinberg, R.A. (1993). Physical interaction of the retinoblastoma protein with human D cyclins. Cell 73, 499–511.

Ewald, J.C. (2018). How yeast coordinates metabolism, growth and division. Current Opinion in Microbiology 45, 1–7.

Ewald, J.C., Kuehne, A., Zamboni, N., and Skotheim, J.M. (2016). The Yeast Cyclin-Dependent Kinase Routes Carbon Fluxes to Fuel Cell Cycle Progression. Molecular Cell 62, 532–545.

Finn, R.S., Martin, M., Rugo, H.S., Jones, S., Im, S.-A., Gelmon, K., Harbeck, N., Lipatov, O.N., Walshe, J.M., Moulder, S., et al. (2016). Palbociclib and Letrozole in Advanced Breast Cancer. https://Doi.org/10.1056/NEJMoa1607303 375, 1925–1936.

Gorges, L.L., Lents, N.H., and Baldassare, J.J. (2008). The extreme COOH terminus of the retinoblastoma tumor suppressor protein pRb is required for phosphorylation on Thr-373 and activation of E2F. Am. J. Physiol., Cell Physiol. 295, C1151–C1160.

Grafström, R.H., Pan, W., and Hoess, R.H. (1999). Defining the substrate specificity of cdk4 kinase–cyclin D1 complex. Carcinogenesis 20, 193–198.

Guiley, K.Z., Liban, T.J., Felthousen, J.G., Ramanan, P., Litovchick, L., and Rubin, S.M. (2015). Structural mechanisms of DREAM complex assembly and regulation. Genes & Development 29, 961–974.

Hafner, M., Mills, C.E., Subramanian, K., Chen, C., Chung, M., Boswell, S.A., Everley, R.A., Walmsley, C.S., Juric, D., and Sorger, P. (2017). Therapeutically advantageous secondary targets of abemaciclib identified by multi-omics profiling of CDK4/6 inhibitors. bioRxiv 211680.

Håland, T.W., Boye, E., Stokke, T., Grallert, B., and Syljuåsen, R.G. (2015). Simultaneous measurement of passage through the restriction point and MCM loading in single cells. Nucleic Acids Research 43, e150–e150.

Hirschi, A., Cecchini, M., Steinhardt, R.C., Schamber, M.R., Dick, F.A., and Rubin, S.M. (2010). An overlapping kinase and phosphatase docking site regulates activity of the retinoblastoma protein. Nat Struct Mol Biol 17, 1051–1057.

Hitomi, M., and Stacey, D.W. (1999). Cyclin D1 production in cycling cells depends on Ras in a cell-cycle-specific manner. Current Biology 9, 1075–S2.

Jaspersen, S.L., Charles, J.F., and Morgan, D.O. (1999). Inhibitory phosphorylation of the APC regulator Hct1 is controlled by the kinase Cdc28 and the phosphatase Cdc14. Current Biology 9, 227–236.

Johnson, D.G., Ohtani, K., and Nevins, J.R. (1994). Autoregulatory control of E2F1 expression in response to positive and negative regulators of cell cycle progression. Genes & Development 8, 1514–1525.

Kato, J., Matsushime, H., Hiebert, S.W., Ewen, M.E., and Sherr, C.J. (1993). Direct binding of cyclin D to the retinoblastoma gene product (pRb) and pRb phosphorylation by the cyclin D-dependent kinase CDK4. Genes & Development 7, 331–342.

Kitagawa, M., Higashi, H., Jung, H.K., Takahashi, I.S., Ikeda, M., Tamai, K., Kato, J., Segawa, K., Yoshida, E., Nishimura, S., et al. (1996). The consensus motif for phosphorylation by cyclin D1-Cdk4 is different from that for phosphorylation by cyclin A/E-Cdk2. The EMBO Journal 15, 7060–7069.

Kõivomägi, M., Valk, E., Venta, R., Iofik, A., Lepiku, M., Morgan, D.O., and Loog, M. (2011). Dynamics of Cdk1 Substrate Specificity during the Cell Cycle. Molecular Cell 42, 610–623.

Kramer, E.R., Scheuringer, N., Podtelejnikov, A.V., Mann, M., and Peters, J.M. (2000). Mitotic regulation of the APC activator proteins CDC20 and CDH1. Mol. Biol. Cell 11, 1555–1569.

Loog, M., and Morgan, D.O. (2005). Cyclin specificity in the phosphorylation of cyclin-dependent kinase substrates. Nature 434, 104–108.

Lundberg, A.S., and Weinberg, R.A. (1998). Functional inactivation of the retinoblastoma protein requires sequential modification by at least two distinct cyclin-cdk complexes. Mol. Cell. Biol. 18, 753–761.

Malumbres, M., and Barbacid, M. (2005). Mammalian cyclin-dependent kinases. Trends in Biochemical Sciences 30, 630–641.

Markey, M.P., Bergseid, J., Bosco, E.E., Stengel, K., Xu, H., Mayhew, C.N., Schwemberger, S.J., Braden, W.A., Jiang, Y., Babcock, G.F., et al. (2007). Loss of the retinoblastoma tumor suppressor: differential action on transcriptional programs related to cell cycle control and immune function. Oncogene 26, 6307–6318.

Matsushime, H., Quelle, D.E., Shurtleff, S.A., Shibuya, M., Sherr, C.J., and Kato, J.Y. (1994). D-type cyclin-dependent kinase activity in mammalian cells. Mol. Cell. Biol. 14, 2066–2076.

Matsuura, I., Denissova, N.G., Wang, G., He, D., Long, J., and Liu, F. (2004). Cyclin-dependent kinases regulate the antiproliferative function of Smads. Nature 430, 226–231.

Mazur, P.K., Herner, A., Mello, S.S., Wirth, M., Hausmann, S., Sánchez-Rivera, F.J., Lofgren, S.M., Kuschma, T., Hahn, S.A., Vangala, D., et al. (2015). Combined inhibition of BET family proteins and histone deacetylases as a potential epigenetics-based therapy for pancreatic ductal adenocarcinoma. Nature Medicine 21, 1163–1171.

Medina, E.M., Turner, J.J., Gordân, R., Skotheim, J.M., and Buchler, N.E. (2016). Punctuated evolution and transitional hybrid network in an ancestral cell cycle of fungi. eLife 5, 120.

Merrick, K.A., Wohlbold, L., Zhang, C., Allen, J.J., Horiuchi, D., Huskey, N.E., Goga, A., Shokat, K.M., and Fisher, R.P. (2011). Switching Cdk2 on or off with small molecules to reveal requirements in human cell proliferation. Molecular Cell 42, 624–636.

Morgan, D.O. (1997). Cyclin-dependent kinases: engines, clocks, and microprocessors. Annu. Rev. Cell Dev. Biol. 13, 261–291.

Morgan, D.O. (2007). The Cell Cycle (New Science Press).

Narasimha, A.M., Kaulich, M., Shapiro, G.S., Choi, Y.J., Sicinski, P., and Dowdy, S.F. (2014). Cyclin D activates the Rb tumor suppressor by mono-phosphorylation. eLife 3, 1068.

Nick Pace, C., and Martin Scholtz, J. (1998). A Helix Propensity Scale Based on Experimental Studies of Peptides and Proteins. Biophysical Journal 75, 422–427.

Pan, W., Cox, S., Hoess, R.H., and Grafström, R.H. (2001). A Cyclin D1/Cyclin-dependent Kinase 4 Binding Site within the C Domain of the Retinoblastoma Protein. Cancer Res 61, 2885–2891.

Pardee, A.B. (1974). A restriction point for control of normal animal cell proliferation. PNAS 71, 1286–1290.

Rubin, S.M., Gall, A.-L., Zheng, N., and Pavletich, N.P. (2005). Structure of the Rb C-Terminal Domain Bound to E2F1-DP1: A Mechanism for Phosphorylation-Induced E2F Release. Cell 123, 1093–1106.

Russo, A.A., Jeffrey, P.D., Patten, A.K., Massagué, J., and Pavletich, N.P. (1996). Crystal structure of the p27Kip1 cyclin-dependent-kinase inibitor bound to the cyclin A–Cdk2 complex. Nature 382, 325–331.

Sack, L.M., Davoli, T., Li, M.Z., Li, Y., Xu, Q., Naxerova, K., Wooten, E.C., Bernardi, R.J., Martin, T.D., Chen, T., et al. (2018). Profound Tissue Specificity in Proliferation Control Underlies Cancer Drivers and Aneuploidy Patterns. Cell.

Salazar-Roa, M., and Malumbres, M. (2017). Fueling the Cell Division Cycle. Trends in Cell Biology 27, 69–81.

Schulman, B.A., Lindstrom, D.L., and Harlow, E. (1998). Substrate recruitment to cyclin-dependent kinase 2 by a multipurpose docking site on cyclin A. Pnas 95, 10453–10458.

Schwarz, C., Johnson, A., Kõivomägi, M., Zatulovskiy, E., Kravitz, C.J., Doncic, A., and Skotheim, J.M. (2018). A Precise Cdk Activity Threshold Determines Passage through the Restriction Point. Molecular Cell 69, 253–264.e255.

Sherr, C.J., and McCormick, F. (2002). The RB and p53 pathways in cancer. Cancer Cell 2, 103–112.

Sherr, C.J., and Roberts, J.M. (2004). Living with or without cyclins and cyclin-dependent kinases. Genes & Development 18, 2699–2711.

Sherr, C.J., Beach, D., and Shapiro, G.I. (2016). Targeting CDK4 and CDK6: From Discovery to Therapy. Cancer Discov 6, 353–367.

Stern, B., and Nurse, P. (1996). A quantitative model for the cdc2 control of S phase and mitosis in fission yeast. Trends Genet. 12, 345–350.

Swaffer, M.P., Jones, A.W., Flynn, H.R., Snijders, A.P., and Nurse, P. (2016). CDK Substrate Phosphorylation and Ordering the Cell Cycle. Cell 167, 1750–1761.e16.

Swaffer, M.P., Jones, A.W., Flynn, H.R., Snijders, A.P., and Nurse, P. (2018). Quantitative Phosphoproteomics Reveals the Signaling Dynamics of Cell-Cycle Kinases in the Fission Yeast Schizosaccharomyces pombe. Cell Rep 24, 503–514.

Takeda, D.Y., Wohlschlegel, J.A., and Dutta, A. (2001). A bipartite substrate recognition motif for cyclin-dependent kinases. J. Biol. Chem. 276, 1993–1997.

The, I., Ruijtenberg, S., Bouchet, B.P., Cristobal, A., Prinsen, M.B.W., van Mourik, T., Koreth, J., Xu, H., Heck, A.J.R., Akhmanova, A., et al. (2015). Rb and FZR1/Cdh1 determine CDK4/6-cyclin D requirement in C. elegans and human cancer cells. Nat Commun 6, 5906.

Wallace, M., and Ball, K.L. (2004). Docking-Dependent Regulation of the Rb Tumor Suppressor Protein by Cdk4. Mol. Cell. Biol. 24, 5606–5619.

Wang, H., Nicolay, B.N., Chick, J.M., Gao, X., Geng, Y., Ren, H., Gao, H., Yang, G., Williams, J.A., Suski, J.M., et al. (2017). The metabolic function of cyclin D3–CDK6 kinase in cancer cell survival. Nature 546, 426–430.

Wohlschlegel, J.A., Dwyer, B.T., Takeda, D.Y., and Dutta, A. (2001). Mutational analysis of the Cy motif from p21 reveals sequence degeneracy and specificity for different cyclin-dependent kinases. Mol. Cell. Biol. 21, 4868–4874.

Yung, Y., Walker, J.L., Roberts, J.M., and Assoian, R.K. (2007). A Skp2 autoinduction loop and restriction point control. J. Cell Biol. 178, 741–747.

Zachariae, W., Schwab, M., Nasmyth, K., and Seufert, W. (1998). Control of cyclin ubiquitination by CDK-regulated binding of Hct1 to the anaphase promoting complex. Science 282, 1721–1724.

Zetterberg, A., and Larsson, O. (1985). Kinetic analysis of regulatory events in G1 leading to proliferation or quiescence of Swiss 3T3 cells. Pnas 82, 5365–5369.

Zhao, G., Chen, Y., Carey, L., and Futcher, B. (2016). Cyclin-Dependent Kinase Co-Ordinates Carbohydrate Metabolism and Cell Cycle in S. cerevisiae. Molecular Cell 62, 546–557.

## References for Methods

Buchan, D.W.A., Minneci, F., Nugent, T.C.O., Bryson, K., and Jones, D.T. (2013). Scalable web services for the PSIPRED Protein Analysis Workbench. Nucleic Acids Research 41, W349–W357.

Chytil, A., Waltner-Law, M., West, R., Friedman, D., Aakre, M., Barker, D., and Law, B. (2004). Construction of a cyclin D1-Cdk2 fusion protein to model the biological functions of cyclin D1-Cdk2 complexes. J. Biol. Chem. 279, 47688–47698.

Cock, P.J.A., Antao, T., Chang, J.T., Chapman, B.A., Cox, C.J., Dalke, A., Friedberg, I., Hamelryck, T., Kauff, F., Wilczynski, B., et al. (2009). Biopython: freely available Python tools for computational molecular biology and bioinformatics. Bioinformatics 25, 1422–1423.

Ding, S., Wu, X., Li, G., Han, M., Zhuang, Y., and Xu, T. (2005). Efficient Transposition of the piggyBac (PB) Transposon in Mammalian Cells and Mice. Cell 122, 473–483.

Finn, R.D., Clements, J., Arndt, W., Miller, B.L., Wheeler, T.J., Schreiber, F., Bateman, A., and Eddy, S.R. (2015). HMMER web server: 2015 update. Nucleic Acids Research 43, W30–W38.

Greenfield, N.J., and Fasman, G.D. (1969). Computed circular dichroism spectra for the evaluation of protein conformation. Biochemistry 8, 4108–4116.

Hagopian, J.C., Kirtley, M.P., Stevenson, L.M., Gergis, R.M., Russo, A.A., Pavletich, N.P., Parsons, S.M., and Lew, J. (2001). Kinetic basis for activation of CDK2/cyclin A by phosphorylation. J. Biol. Chem. 276, 275–280.

Katoh, K., Rozewicki, J., and Yamada, K.D. (2017). MAFFT online service: multiple sequence alignment, interactive sequence choice and visualization. Brief. Bioinformatics.

McCusker, D., Denison, C., Anderson, S., Egelhofer, T.A., 3rd, J.R.Y., Gygi, S.P., and Kellogg, D.R. (2007). Cdk1 coordinates cell-surface growth with the cell cycle. Nature Cell Biology 2007 9:5 9, 506–515.

Price, M.N., Dehal, P.S., and Arkin, A.P. (2009). FastTree: computing large minimum evolution trees with profiles instead of a distance matrix. Mol. Biol. Evol. 26, 1641–1650.

Rao, R.N., Stamm, N.B., Otto, K., Kovacevic, S., Watkins, S.A., Rutherford, P., Lemke, S., Cocke, K., Beckmann, R.P., Houck, K., et al. (1999). Conditional transformation of rat embryo fibroblast cells by a *cyclin D1-cdk4* fusion gene. Oncogene 18, 6343–6356.

Shariati, A., Dominguez, A., Wernig, M., Qi, S., and Skotheim, J. (2018). Reversible inhibition of specific transcription factor-DNA interactions using CRISPR. bioRxiv 282681.

